# The post-transcriptional regulation of TFs in immature motoneurons shapes the axon-muscle connectome

**DOI:** 10.1101/2021.10.22.465474

**Authors:** Wenyue Guan, Stéphanie Bellemin, Mathilde Bouchet, Lalanti Venkatasubramanian, Camille Guillermin, Anne Laurençon, Kabir Chérif, Aurélien Darmas, Christophe Godin, Séverine Urdy, Richard S. Mann, Jonathan Enriquez

**Affiliations:** Institut de Génomique Fonctionnelle de Lyon, ENS de Lyon, CNRS, Univ Lyon 1, 46 Allée d’Italie, 69364 Lyon Cedex 07, France; Departments of Biochemistry and Molecular Biophysics, and Neuroscience, Mortimer B. Zuckerman Mind Brain Behavior Institute, Columbia University, New York, NY 10027, USA; Departments of Pathology and Cell Biology, Neuroscience and Ophthalmology, Mortimer B. Zuckerman Mind Brain Behavior Institute, Columbia University, New York, NY 10027, USA; Laboratoire Reproduction et Développement des Plantes, Univ Lyon, ENS de Lyon, UCB Lyon 1, CNRS, INRA, Lyon, France

**Author notes:** Contributed equally to this work (Stéphanie Bellemin, Mathilde Bouchet, Lalanti Venkatasubramanian). Correspondance (J.E.).

## Abstract

Temporal factors expressed sequentially in neural stem cells, such as RNA binding proteins (RBPs) or transcription factors (TFs), are key elements in the generation of neuronal diversity. The molecular mechanism underlying how the temporal identity of stem cells is decoded into their progeny to generate neuronal diversity is largely unknown. Here, we used genetic and new computational tools to study with precision the unique fates of the progeny of a stem cell producing 29 morphologically distinct leg motoneurons (MNs) in *Drosophila*. We identified 40 TFs expressed in this MN lineage, 15 of which are expressed in a combinatorial manner in immature MNs just before their morphological differentiation. By following TF expression patterns at an earlier developmental stages, we discovered 19 combinatorial codes of TFs that were progressively established in immature MNs as a function of their birth order. The comparison of the RNA and protein expression profiles of 6 TFs revealed that post-transcriptional regulation plays an essential role in shaping these TF codes. We found that the two known RBPs, Imp and Syp, expressed sequentially in neuronal stem cells, are upstream regulators of the TF codes. Both RBPs are key players in the construction of axon-muscle connectome through the post-transcriptional regulation of 5 of the 6 TFs examined. By deciphering the function of Imp in the immature MNs with respect to the stem cell of the same lineage, we propose a model where RBPs shape the morphological fates of MNs through post-transcriptional regulation of TF codes in immature MNs. Taken together, our study reveals that immature MNs are plastic cells that have the potential to acquire many morphological fates. The molecular basis of MN plasticity originates in the broad expression of different TF mRNA, that are post-transcriptionally shaped into TF codes by Imp and Syp, and potentially by other RBPs that remain to be discovered, to determine their morphological fates.

## INTRODUCTION

Most, if not all animal behaviors, such as eating, walking, mating or playing the piano, requires coordinated and precise movements. As Michel De Montaigne said: ‘our life is only movement’. Locomotion is a stereotyped behavior used by animals to find food, mates or escape from predators. The rhythmic and precise pattern of locomotion is linked to the coordinated contraction of muscles which are innervated by a complex wiring of motor neuron axons controlling the timing and the intensity of muscle activity. The architecture of this precise cabling is stereotyped in animals of the same species from invertebrates to vertebrates and is built during the morphological differentiation of motoneurons (MNs). One central challenge is to decipher how stem cells can generate a huge diversity of MN morphologies. In our study, we used a combination of genetic and computational approaches to understand how in *Drosophila* a single stem cell gives rise to 29 morphologically unique motor neurons.

Transcription factors (TF) are central regulators of the morphological specification of neurons, including MNs. In vertebrates, during development, Hox6 and Hox10 paralogs at the brachial and lumbar levels of the spinal cord, respectively, distinguish MNs that target leg muscles from those that target the body wall muscles. Subsequently, limb-targeting MNs are further refined into pools, where all MNs in a single pool target the same muscle. Each pool is molecularly defined by the expression of pool-specific TFs (Dasen and Jessell, 2009; Philippidou and Dasen, 2013). In *Drosophila* embryos, subclasses of MNs innervating the larva body-wall muscles are also morphologically specified, in terms of axonal targeting, by unique combinations of TFs: *evenskipped* (eve) and *grain* are expressed in six MNs that target dorsal body-wall muscles (Fujioka et al., 2003; Garces and Thor, 2006; Landgraf et al., 1999), and *Hb9, Nkx6, Islet, Lim3* and *Olig2* are required in ventral-targeting MNs (Broihier and Skeath, 2002; Broihier et al., 2004; Certel and Thor, 2004; Oyallon et al., 2012; Thor and Thomas, 1997; Thor et al., 1999). In adult *Drosophila*, the molecular specification of MN morphologies in the leg, which can be defined by a specific dendritic arbor and axonal targeting, seems to be controlled at the single-cell level by a combination of morphology-specifying TFs (mTFs) expressed during larval and pupal development. Seven out of the ~50 MNs innervating limb appendages differentially express unique combinatorial codes of five mTFs (mTFs, *Pb, ems, toy, zfh1, zfh2*) that can account for most of their morphological diversity. Strikingly, the reprograming of these mTF codes is sufficient to lead to switches in morphology (Enriquez et al., 2015).

Immature neurons expressing the TF codes are generated by dedicated stem cells. These stem cells are regulated in space and time to generate the enormous amount of morphological diversity at the right time and in the right place (Kohwi and Doe, 2013; Sagner and Briscoe, 2019). In *Drosophila*, neurons are generated by neuroblasts (NBs), specialized stem cells dedicated to the generation of neurons and glia (Doe and Skeath, 1996; Prokop and Technau, 1991; Truman and Bate, 1988). As they divide, NBs express a temporal sequence of transcription factors (tTFs) that contributes to the generation of neuronal diversity. In the embryonic ventral nerve cord (VNC; and analogue of the human spinal chord) most NBs express a sequence of five tTFs (Hunchback, Krüppel, Pdm1/Pdm2, Castor and Grainyhead) (Isshiki et al., 2001; Li et al., 2013a), whereas in medulla NBs of the visual system and intermediate neural progenitors of the *Drosophila* larval brain, a different series of tTFs have been described (Bayraktar and Doe, 2013; Konstantinides et al., 2021; Li et al., 2013b). In vertebrates, neural stem cells, e.g. in the cerebral cortex, retina and spinal cord, use analogous strategies suggesting that the regulatory logic of tTFs changing the identity of stem cells through development is evolutionarily conserved (Alsiö et al., 2013; Delile et al., 2019; Elliott et al., 2008; Jacob et al., 2008; Mattar et al., 2015; Okano and Temple, 2009).

In *Drosophila*, neuronal diversity can be generated by NBs as they age via a second mechanism involving two RNA-binding proteins (RBPs), IGF-II mRNA-binding protein (Imp) and Syncrip (Syp). These RBPs are expressed sequentially in brain NBs from high-to-low and low-to-high, respectively, over time (Liu et al., 2015; Syed et al., 2017). In the NBs of the mushroom body, a structure in the brain processing olfactory inputs, the two RBPs regulate the translation of the TF Chinmo, which in turn controls the temporal identity of the mushroom-body NB progeny in a concentration-dependent manner. This gradient strategy allows the generation of many neurons of the same type, as observed in mammalian progenitors (Liu et al., 2015; Zhu et al., 2006).

In *Drosophila*, most of the MNs innervating leg muscles are derived from around 10 lineages in each ganglion (Baek and Mann, 2009; Brierley et al., 2009, 2012; Maniates-Selvin et al., 2020). These lineages produce ~50 immature MNs (iMNs) during larval stages that differentiate morphologically during metamorphosis to acquire their specific dendritic arbors and axonal targeting. Here we used a lineage called Lin A/15, which produces glial cells and MNs (Baek et al., 2013; Enriquez et al., 2018), to understand how the generation of different neuronal morphologies is molecularly controlled during development. We choose this lineage for multiple reasons. First, the Lin A/15 NB produces two thirds of the adult limb MNs (Baek and Mann, 2009). Second, with each division, the Lin A/15 NB generates a MN that eventually establishes a unique and invariable axonal connection with distinct leg muscles. In other words, each MN morphology is unique and linked to its birth order (Baek and Mann, 2009). The complex architecture of the neuromuscular system, which executes locomotion in adult *Drosophila* (Azevedo et al., 2020) is made possible by the generation of a stereotyped axon-muscle connectome, which must be shaped during development by a sophisticated molecular logic. To gain insight into this logic, we screened for TFs expressed in iMNs just before their morphological differentiation.

Here, we report the discovery of 40 TFs expressed in immature Lin A/15 MNs, 15 of which are differentially expressed in iMNs just before their morphological differentiation. By combining (i) a new computational method to correlate birth order and the position of iMNs relative to that of the NB with (ii) multiple immunostaining, we characterized 19 different codes of TFs expressed in the 29 iMNs as a function of their birth order. We demonstrated that the TF code is not established at the birth of the MN but is gradually shaped during development. By comparing the expression profile of 6 mRNAs with their corresponding proteins, we revealed that post-transcriptional regulation of TFs plays a key role in shaping the TF code. We then analyzed the function of two known RBPs, Imp and Syp, in shaping the code in iMNs. We first demonstrated that in earlier born MNs, Syp protein was undetectable and the level of Imp was high, whereas in the later born MNs, the level of Syp protein was high and Imp was low. The functional analysis of both proteins revealed that they are key players on shaping the axon-muscle connectome by regulating the translation of at least 5 TFs. Finally, we deciphered the function Imp in iMNs versus the NB and revealed that Imp controls the fate of the Lin A/15 progenies directly in iMNs. Together, these experiments allow us to propose a model where RBPs control the fate of iMNs by post-transcriptionally sculpting a complex set of TF codes.

## RESULTS

### Lin A, a model to study how the MN-muscle connectome is built during development

Previous studies have characterized Lin A/15 MNs by MN-driven GFP labelling, whereby the presumptive muscle target is identified through the localization of the terminal branches of the individual MN in the leg (Baek and Mann, 2009; Brierley et al., 2009, 2012).

Here, we extend these earlier studies by driving the expression of RFP in all leg muscles (*Mhc-RFP*), in addition to driving the expression of GFP in 28 Lin A/15 MNs with GFP (*VGlut>GFP*, MARCM; **Fig. 1D1-D2; see note S1 the comparison with other studies)**. Lin A/15 MNs innervate 9 out of the 14 leg muscles: all 5 muscles in the tibia (tarm1/2, tadm, talm, ltm1), three muscles in the femur (tirm, tidm, ltm2) and one muscle in the trochanter (ferm). The first-born MN from Lin A/15, which cannot be visualized with the “one-spot” MARCM technique **(see note S2)**, innervates a body wall muscle and can be genetically labeled with GFP by using a Lin A/15 tracing system **(Fig. 1A-C, see STAR methods)**. Based on this and previous studies, the tight correlation between the birth order of Lin A/15 MNs and their muscle targets can be schematically summarized (**Fig.1E, note S2)**. Notably, although clusters of MNs born in the same time window frequently target the same muscle, each MN targets a specific fiber based on its birth order, which make each one unique in terms of axonal targeting (Baek and Mann, 2009). The precise characterization of the development of Lin A /15 MNs and the available genetic tools provide the basis to understand the correlation between MN birth order and the specification of morphological identity.

**Figure 1.**
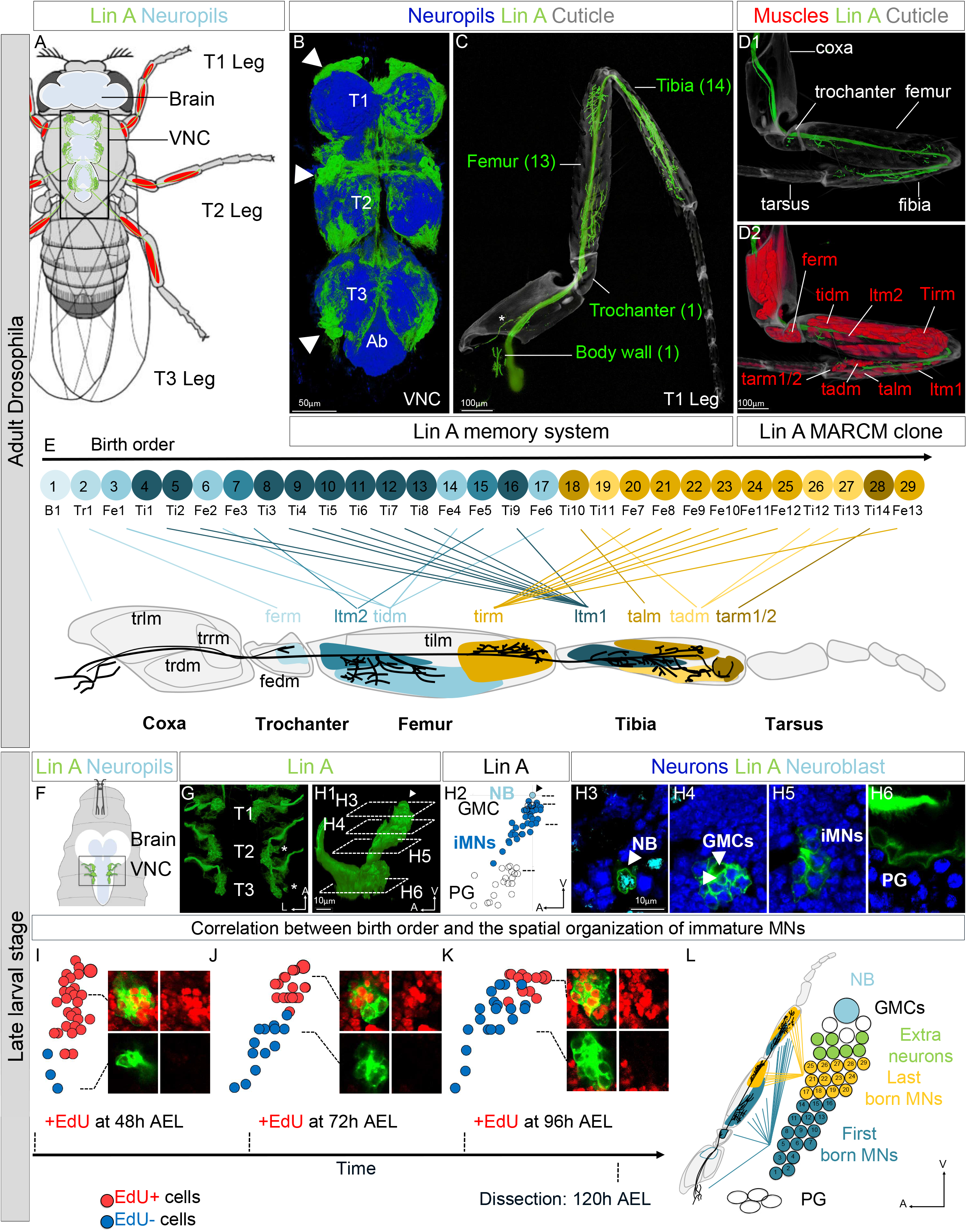
Lin A/15 as model to study how the axon-muscle connectome is built. **(A)** Drawing of an adult fly showing the position of the CNS (white: cortex, blue: neuropiles) and Lin A/15 leg MNs (green cell bodies and dendrites in the VNC and axons in the legs). Black box indicates the VNC imaged in **B**. **(B)** 3D reconstruction of confocal sections of a VNC where the six Lin A/15 were genetically labeled with mCD8::GFP (green) and Neuropil immunostained with anti-BRP (blue). **(C)** 3D reconstruction of confocal sections of a T1 leg where Lin A/15 was genetically labeled as in (**B**) with mCD8::GFP (green). (**D1-D2)** 3D reconstruction of confocal sections of a T1 leg containing a Lin A/15 MARCM clone expressing mCD8:GFP under the control of *VGlut-Gal4* (**D1**-**D2**), and where all muscles were labeled with *Mhc* (Myosin heavy chain)-RFP (**D2**; red). The segments innervated by Lin A/15 are indicated in **(C)**, the leg muscles innervated by Lin A/15 are indicated in **(D2)**, all the leg segment names are indicated in **(D1)**. (**E)** Schematic showing the link between birth order and muscle targeting, modified from (Baek and Mann, 2009) (see Supplemental Information). Top: schematic of the cell body of the 29 LinA/15 MNs. The numbers inside indicate their birth order. The abbreviations below the schematic of the cell body indicate the name of the MNs based on their birth order and the segment they target, nomenclature from (Baek and Mann, 2009). Bottom: schematic of a T1 leg innervated by Lin A/15, the muscles and the name of the muscles innervated by Lin A/15 axons are color-coded based on their innervation. The line between the schematic of the cell body and leg muscles indicates the relationships between MN birth order and muscle innervation. Nomenclature of the muscle is based on (Soler et al., 2004): tr: trochanter, fe: femur, ti: tibia, ta: tarsus, r: reductor, d: depressor, l:levator, m:muscle. **(F)** Drawing of the anterior region of a third instar larva showing the position of the CNS (white: cortex blue: neuropil) and immature Lin A/15 MNs (green). **(G-H2)** 3D reconstruction of confocal sections of six **(G)** and one **(H)** rtight thoracic hemisegment 2 (T2R) Lin A/15 in a third instar larva genetically labeled as in **(B)** with myr::GFP (green). Ventral **(G)** and lateral **(H)** view; axes: Anterior (A), Lateral (L), Ventral (V). **(H1)** Plot of the relative position of each Lin A cell in **(H1)** from a lateral perspective. Lin A proliferative glia (PG) are in white, Lin A/15 iMNs are in blue, Lin A/15 GMCs are in white and Lin A NB is in Cyan. The NB is positioned at the origin of the graph. **(H3-H6)** Confocal sections of the Lin A/15 in **(H1-H2)** immunostained with anti-Elav (blue) and Dpn (cyan). **(I-L)** Plots of the relative position of each Lin A/15 cell from a lateral perspective in third instar larvae fed with Edu at different time points. Edu+ cells are in red, Edu-cells are in blue. The horizontal axis indicates the time point of Edu feeding. On the right of each graph, confocal sections of the Edu- and Edu+ cells. **(L)** Schematic of the cell bodies of Lin A/15 in a third instar larva and of an adult leg. The schematic shows the correlation between the position of the immature MNs and their muscle targeting in adult leg.

### Correlation between birth order and the spatial organization of iMNs

The late third instar is a key developmental stage between the end of MN production and the establishment of stereotyped axon-muscle connections (Baek et al., 2013; Venkatasubramanian et al., 2019). We thus chose this stage as an entry point to understand how MN diversity is generated during development.

We first determined how the spatial organization of iMNs is related to birth order (**Fig. 1F-K)**. In late third instar larva (L3), the NB has a ventral location, whereas the iMNs are located more dorsally and anteriorly **(Fig 1G-H6)**. When larvae are fed with the DNA-label EdU at early time points, the last-born MNs (EdU+) are located ventrally and anteriorly near to the NB, whereas older MNs (EdU-) are located dorsally and anteriorly distant from the NB. Moreover, EdU+ iMNs and EdU-iMNs are barely intermingled **(Fig 1I-K)**. This demonstrates that up to at least the end of the third instar, iMNs maintain distinct spatial positions that are dictated by their birth dates. Importantly for our study, the most ventral MNs, which are the last born iMNs, apoptose during the pupal stages (Wenyue *et al., unpublished*), whereas the first-born 29 iMNs morphologically differentiate during metamorphosis **(Fig. 1L)**. This clear relationship between birth order and the spatial organization of iMNs was then used to understand how iMNs are morphologically specified by TF.

### A complex combination of TFs in iMNs prefigure morphological diversity of LinA/15 MNs

To identify the TFs expressed in Lin A/15 iMNs in L3 VNC that could control the stereotypic wiring of muscle innervation, we performed an immunostaining screen of GFP-expressing iMNs with a collection of around 250 antibodies directed against different TFs, covering ~35 % of all TFs coded by the *Drosophila* genome, (**see STAR methods)**. Lin A/15 NBs and postmitotic MNs were immunolocalized with anti-Dpn and anti-Elav respectively, to distinguish them from the GMCs (Dpn-, Elav- and ventrally localized) and the proliferating glia (Dpn-, Elav- and dorsally localized) **(Figure 1H1-H2, Fig 2A-G2** illustrates the example of the TF, Jim). In total, 43 TFs were identified in Lin A/15, with 16 TFs expressed in clusters of iMNs **(Fig. S1)**. We then focused our study on the 16 TFs expressed in subpopulation of MNs since their differential expressions could be at the origin of the incredible diversity of MN morphologies generated at each division by Lin A/15 NB.

**Figure 2.**
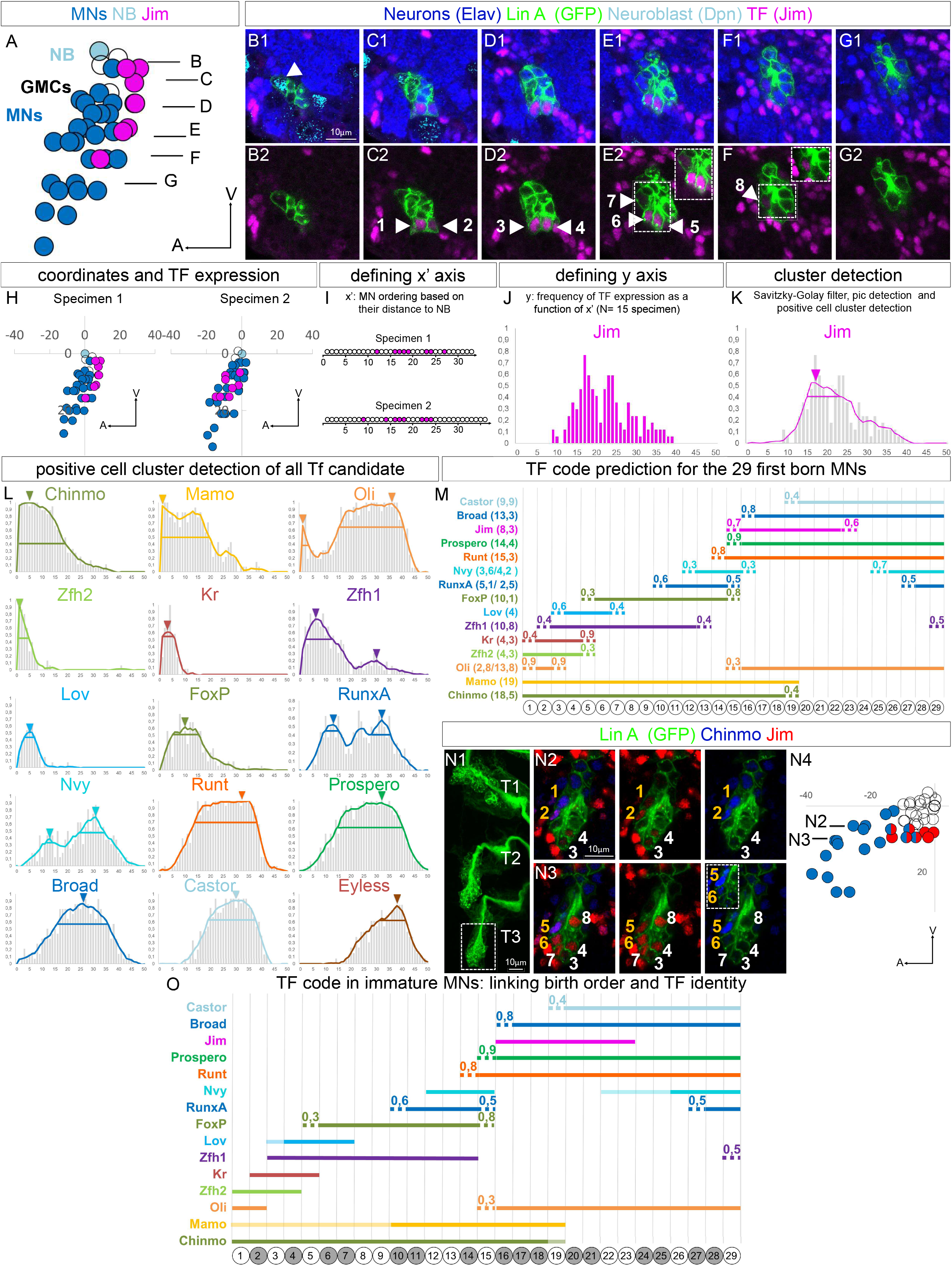
Correlation between birth order and TF codes. **(A)** Plot of the relative position of each Lin A/15 cell from a lateral perspective in a third instar larva immuno-stained with anti-Jim, anti-Elav, anti-Dpn and where Lin A/15 is genetically labeled with myr:GFP. GMCs are in white, Jim-iMNs are in blue, Jim+ iMNs are in purple and Lin A/15 NB is in Cyan. **(B1-G2)** Confocal sections of the Lin A/15 in **(A)**. Anti-Elav (blue), GFP (green), anti-Dpn (cyan), anti-Jim (purple). **(see also Fig S1 for the expression pattern of all TFs)** **(H)** On the left (specimen 1) same graph as in **(A)** but where the spatial axis is represented (units are in micrometer). On the right similar graph from another specimen. **(I)** MN ordering on a x’ axis according to their relative distance from the Lin A NB. The iMNs expressing Jim are color-coded in purple. **(J)** Frequency of Jim expression as a function of x for 15 specimens. **(J)** Frequency of Jim expression as a function of x after applying a Savitzky-Golay filter to smoothen the distribution in (J). The arrow indicates the peak of the Jim+ cell cluster **(see STAR method)**. The horizontal bar indicates the Jim+ cell cluster detected with the PCCD method **(see STAR method)**. **(L)** Frequency of 15 TF expressions as a function of x after applying a Savitzky-Golay filter to smoothen the distribution. The arrows indicate the peak of the TF+ cell clusters **(see STAR method)**. The horizontal bars indicate the TF+ cell clusters detected with the PCCD method **(see STAR method)**. **(M)** schematic of the code expressed in each iMN predicted by the PCCD method in a third instar larva **(see STAR method)**. Bottom: schematic of the cell body of the 29 Lin A/15 MNs. The number inside indicate their birth order. Bottom: the horizontal bars indicate the TF+ cell clusters detected with the PCCD method for 15 TFs. The dotted lines indicate the coverage index at the border of all cell clusters **(see STAR method)**. The numbers in parenthesis indicate the length of the TF positive cell cluster **(see STAR method)**. **(N1-N3)** Max projection **(N1)** of the three left thoracic hemisegments (T1-T3) and two confocal **(N2-N3)** sections of the doted box in **(N1)** in a third instar larva CNS where of Lin A/15 was genetically labeled with myr::GFP (green) and immunostained with anti-Jim (red), anti-Chinmo (blue). The numbers indicated the Jim+ iMNs, the numbers in orange indicate the iMNs expressing both Jim and Chinmo. The position of the confocal sections is indicated in **(N4)**. **(N4)** Plots of the relative position of the Lin A/15 cells from a lateral perspective corresponding to the Lin A/15 boxed in **(N1)**, blue: Chinmo+ MNs, red: Jim+ MNs. Note: one of the Jim+ cell expressed a very low level of Chinmo. (**see Fig. S1 for all co-stainings**). **(O)** Same schematic as in **(M)** after validation and small corrections of the PCCD method by performing co-staining **(see STAR method, supplemental information and Fig S2 for all co-staining)**. Note the TF gradient expressions of Castor, Broad, Runt, Prospero, and Oliwas not taken into consideration when assigning the codes. The weak expression of Chinmo, Nvy and Mamo is indicated in lighter colors. The color change from white to gray each time a new TF code is detected.

We developed a method to assign the correct combination of TFs expressed in each immature Lin A/15 MN as a function of their birth-order. In late L3, the attribution of the accurate combination of TF to each of the 29 iMNs is extremely challenging since iMNs cannot be identified by their morphologies as in the adult. We have therefore developed a method based on the correlation between the relative position of iMNs from Lin A/15 NB and their birth order **(see STAR methods)**. We named this method the Positive Cell Cluster Detection (PCCD) method. In this method, (i) x, y, and z coordinates and on/off expression of a given TF were assigned to each Lin A/15 iMN for at least 15 Lin A/15 samples per immunostaining (**Fig 2H** illustrates two Lin A/15 samples with anti-Jim immunostaining, **Fig. S1** illustrates all TF expression). (ii) With these coordinates, each Lin A/15 iMN is ranked **along an x’ axis** according to its relative distance from the NB, such that the lowest rank (1) corresponds to the greatest distance from the NB **(Fig. 2I,** illustrates the MN ranking of two samples of anti-Jim immunostaining). (iii) The frequency of TF expression is calculated as a function of x’ (**Fig. 2J** illustrated with anti-Jim immunostaining). (iv) The distributions of frequencies are smoothened by application of a Savitzky-Golay filter (polynomial smoothing). (v) The peak of each distribution is identified and the positive cell cluster of iMNs associated with each peak is assigned (**Fig. 2K** illustrates the identification of the iMN cluster expressing Jim, **Fig. 2L** shows similar cluster predictions for all other TFs). We schematized our results for the 29 first-born MNs on a birth order axis in order to predict the combination of TFs expressed in each given MN since there is a correlation between MN birth order and their relative distance to the NB (x’ axis). PCCD reveals that 15 out of the 16 differentially expressed TFs are expressed in combination in the 29 first-born Lin A/15 iMNs **(Fig. 2M)**. To validate and refine this analysis, 12 co-immunostainings for different TFs were performed **(Fig. 2 N1-N4 illustrates the example of Chinmo and Jim co-staining, Fig.S2 show all co-staining)**. The result of the co-stainings **(see supplemental note S3)** reveals that PCCD is very accurate because in most cases none or only one cell correction was required **(see supplemental procedure S1, compare Fig. 2M vs Fig. 2O)**. By combining the cluster detection method with co-staining experiments, we assigned 18 different TF codes for 29 MNs. These results indicate that the diversity and stereotypy of MN morphologies correlates with a differential expression of TF codes linked to MN birth-order. The next step was to discover how the TF code is established during development.

### TF-code diversity is progressively established in iMNs during development

We then determined how the expression pattern of the 16 differentially-expressed-TFs is established by analyzing their expression from the end of L2, when the NB begins to proliferate, until the end of L3.

We classified Lin A/15 TFs into three categories based on their spatial and temporal expression dynamics. The first category encompasses 10 TFs: Zfh1, Zfh2, Kr, Lov, Oli, RunXA, Jim, Castor, Prospero and FoxP, whose expression starts in the GMC or postmitotic neurons (**Fig 3A1-H, Fig S3,** note that Prospero and FoxP were placed in this category even though they were also detected in the NB cytoplasm). The expression dynamics of the TFs in this category were further subdivided into 2 subcategories based on their temporal expression dynamics. Three TFs, Oli, Castor and Prospero (Category 1.1), are expressed in all new-born iMNs. However, their expression is only maintained in a subpopulation of iMNs (**Fig 3E1-H)**. The other sub-category (Category 1.2) was composed of TFs expressed and maintained by subpopulations of iMNs since their birth (Lov, Kr, RunXA, Zfh1 and Zfh2), or whose expression turns on from birth after a delay (Jim and FoxP) (**Fig 3A1-D)**. The second category is composed by Runt, Broad and Nvy, which are TFs expressed continuously in the Lin A/15 NB from late L2 until late L3. All new born iMNs expressed the second category of TFs but their expression is progressively and differentially maintained in different groups of iMNs (**Fig 3I1-L)**. Finally, the third category is composed by Chinmo and Mamo, which are the only TF having a temporal expression in the NB (**Fig 3M1-P)**. As described for other lineages (Dillard et al., 2018; Maurange et al., 2008; Syed et al., 2017), Chinmo is expressed in the NB in early stages and only maintained in the first born MNs. However, the expression of Mamo in the NB is completely uncoupled from its expression in iMNs **(Fig S3)**.

**Figure 3.**
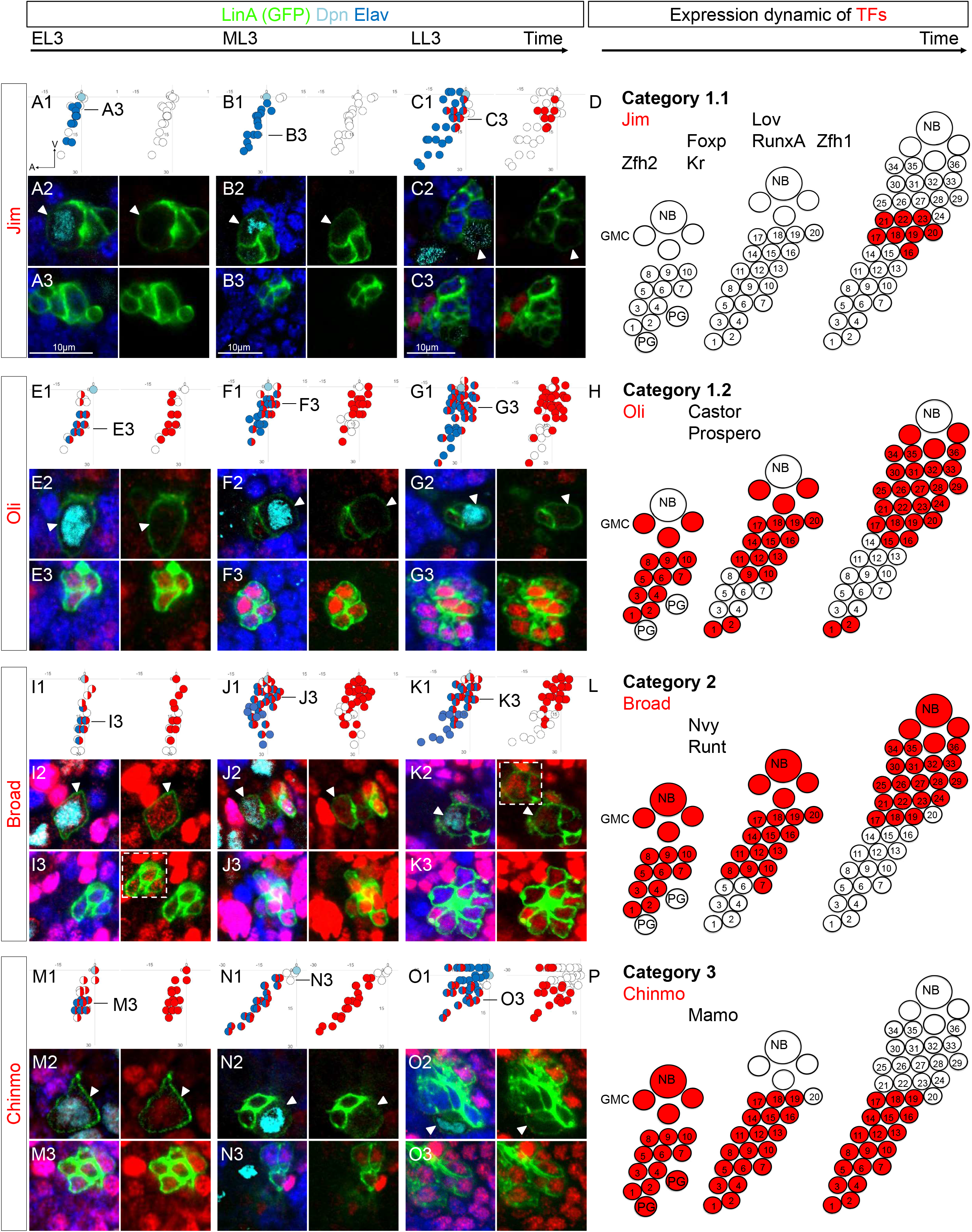
TF expression is progressively shaped in postmitotic neurons during development. **(A-C, E-G, I-K, M-O)** Plot of the relative position of each Lin A/15 cell from a lateral perspective **(A1, B1, C1, D1, E1, F1, G, H1, I1, J1, K1, L1, M1, N1, O1)** and confocal sections **(A2-A3, B2-B3, C2-C3, D2-D3, E2-E3, F2-F3, G2-G3, H2-H3, I3-I3, J2-J3, K2-K3, L2-L3, M2-M3, N2-N3, O2-O3)** showing the expression of Jim **(A1-C3)**, Oli **(E1-G3)**, Br **(I1-K3)** and Chinmo **(M1-O3)** (red) in Lin A/15 genetically labeled with GFP (green) and immunostained with anti-Dpn (cyan), anti-Elav (blue) throughout development. Arrowheads indicate the NB. The developmental time points are indicated on top (EL3/ML3/LL3: Early/Mid/Late third-instar larvae). The expression pattern of each given TF was evaluated at 7 different time points: EL3 (64-67 hours AEL; 70-73 hours AEL; 77-80 hours AEL), ML3 (88-91 hours AEL; 94-97 hours AEL; 101-104 hours AEL) and LL3 (120 hours AEL). representative times points are chosen for EL3/ML3/LL3 depending of the TFs.**(A1, B1, C1, D1, E1, F1, G, H1, I1, J1, K1, L1, M1, N1, O1,)** Axes: Anterior (A), Ventral (V). Left panel: Lin A/15 NB is in cyan, Lin A/15 GMCs (near NB) and proliferative glia (intermingled with MNs at early time points) are in white, Lin A/15 immature MNs are in blue, the cell expressing a TF are indicated with red. Right panel: only the TFs-expressing cells are color-coded in red. The black lines indicate the positions of the confocal section in **(A3, B3, C3, E3, F3, G3, I3, J3, K3, M3, N3, O3)**. The confocal section in **(A2, B2, C2, E2, F2, G2, I2, J2, K2, M2, N2, O2)** are always thought the NB. **(D, H, L, P)** Schematic of the expression of Jim **(D)**, Oli **(H)**, Br **(L)** and Chinmo **(P)** in Lin A/15 during larval stages. Jim/Oli/Br/Chinmo-expressing cells are indicated in red. 1-29 numbers the MNs according to their birth order, with 1 indicating the first-born MN. The TFs in the same categories are listed on top left of each schema. **(See Figure S3 for all TF stainings).** NB: neuroblast; GMC: ganglion mother cells; PG: proliferating glia.

Together, these results showed that the TF code was not established in the NB at each division and transferred to the neuronal progeny, but was gradually established in iMNs by a *de novo* expression of TFs in GMCs and post-mitotic neurons (Category 1), and by the selective maintenance and/or repression of TFs in iMNs (Categories 1, 2 and 3). These expression dynamics suggest the existence of upstream regulators shaping the TF codes by maintenance and/or repressive mechanisms.

### Post-transcriptional regulation governs the establishment of TF codes

In view of the complex dynamics of TF protein expression, we wondered to what extent it could be shaped by post-transcriptional regulation. To address this, we analyzed the RNA expression pattern of 6 TFs selected as representatives of the protein-expression profiles categorized above: *jim* and *Oli* (Category 1), *broad* and *nvy* (Category 2) and *mamo* and *chinmo* (Category 3).

We performed smFISH and developed a pipeline to precisely quantify the number of RNA spots per Lin A/15 iMN in the third instar larva. This pipeline combined 3D segmentation of all Lin A/15 iMNs orientated by membrane-GFP labeling (Machado et al., 2019) and a computational quantification of the number of RNA-positive spots in each Lin A/15 cell (Raj et al., 2008)(**Fig 4 A-I and Fig S4)**. This analysis showed that the RNAs of all six TFs were present in more iMNs than the respective proteins **(Fig. 4 A-I vs Fig. 2)**. The average number of RNA dots detected in each iMN (N) as a function the iMN’s relative distance to the NB was calculated for ~4 to 6 samples per RNA type. Averages were also calculated for clusters of contiguous iMNs (N^x’-x”^), where x’ and x” where the first and last iMN in the selected cluster. For example, *jim* RNA was expressed at higher levels in ventral iMNs proximal to the NB (N^21-29^= 52 and N^11-20^= 54) than the dorsal iMNs (N^1-10^= 27) and was barely detectable in the NB (N^NB^= 9) **(Fig. 4 A-E2)**. The cells close to the NB, which express a very low level of *Jim*, were probably GMCs or apoptotic siblings from asymmetric divisions **(Fig. 4D)**. For *chinmo* mRNA, which is known to be expressed in late NB and its progeny in the VNC (Dillard et al., 2018), it was expressed in all iMNs at a similar level (N^1-10^=31, N^21-29^=43, and N^11-20^=36,) and was strongly expressed in the NB (N^NB^= 161) **(Fig. 4D)**. The RNA of *mamo, broad, nvy* and *Oli* were also expressed ubiquitously or in most iMNs **(Fig. 4G-I and Fig. S4).**

**Figure 4.**
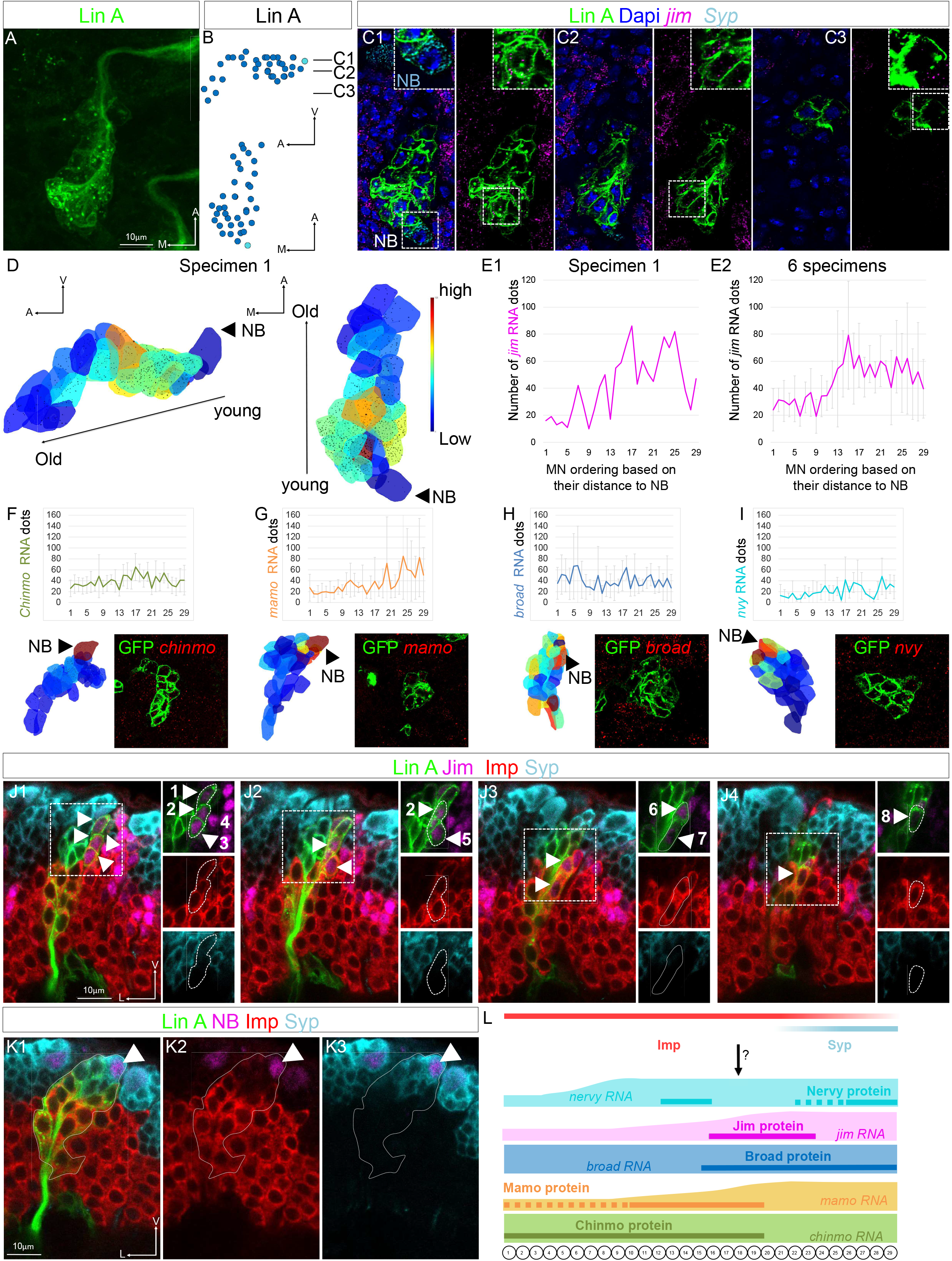
RNA expression of Lin A/15 TFs and of two Imp and Syp RBPs. **(A-C3)** Lin A/15 in a hemisegment T2 (right side). **(A)** Max projection of confocal images where Lin A/15 was genetically labeled with myr::GFP (green). **(B)** Plots of the relative position of each Lin A cell in **(H)** from lateral (top) and ventral (bottom) perspectives. Axes: Anterior (A), Medial (M), Ventral (V). Lin A/15 iMNs and GMCs are both in blue and cannot be distinguished; Lin A NB is in Cyan. **(C1-C2)** Confocal sections of the Lin A/15 in **(A)**. All cells were labeled with dapi (blue); *jim* (purple) and *Syp* (cyan) mRNA. Note: *syp* smFISH also detected the NB. The boxed regions in **(C1-C3)** are enlarged at the top-right region of each panel. Note 1: In **(C1)** the boxed region shows the syp+ NB. Note 2: in (**C**), the intensity of the GFP and smFISH signals was numerically enhanced on the enlarged boxed region to highlight the weak signal. The reduction of the signal intensity in dorsal cells was due to the weak health of the tissue. **(D)** 3D reconstruction of the segmented Lin A/15 cells on **(A)** (**see STAR method)**. Left: lateral view, right: ventral view. Axes: Anterior (A), Medial (L), Ventral (V). Each cell is color-coded based on *jim* RNA relative expression from low (blue) to high (brown). The *jim* mRNA spots are represented by spheres (**see STAR method)**. **(E1-E2)** Plot of the expression level **(E1)** of the *jim* mRNA in the Lin A/15 MNs in **(A)** and plot of the average expression **(E2)** level of the *jim* RNA in 6 Lin A/15 samples, as a function of their relative distance to the NB. Note: only the 29 MNs most distant to NB are represented. **(F-I)** Top: graphs of the average expression level of each RNA as a function of their relative distance from the NB for a minimum of N=4 (N, Number of Lin A/15). Note: only the 29 MNs most distant to NB are represented. (**See Fig. S4 for *Oli*** *m***RNA expression).** Bottom left: 3D reconstruction (as in D) of segmented Lin A/15 cells where *chinmo* **(F)**, *mamo* **(G)**, broad**(H)** and *nvy* **(I)** RNA are detected by smFISH. Bottom left: one confocal section. *mamo* expression: N^1-10^=21, N^21-29^= 24, N^11-20^=51, N^NB^= 89, average <u>N</u>umber for 6 samples of *mamo* RNA spots detected in iMNs at position 1-to 10, 11 to 20, 21 to 29_and in the NB. *broad* expression: N^1-10^=38, N^21-29^= 38, N^11-20^=36, N^NB^= 62, average <u>N</u>umber for 5 samples of *broad* RNA spots detected in t iMNs). *Nvy* expression: N^1-10^=8, N^21-29^= 20, N^11-20^=21, N^NB^= 33, average <u>N</u>umber for 4 samples of *nvy* RNA spots detected in iMNs) **(J1-J4)** Confocal sections of a T2 left Lin A/15 genetically labeled with myr::GFP (green) and immunostained with anti-Jim (purple), Syp (cyan) and Imp (red). The boxed regions and the arrowhead indicate the Jim+ MNs. Note: the CNS was mounted laterally to visualize on the same section old vs young MNs (**see STAR method)**. **(K1-3)** A confocal section of a T2 left Lin A/15 genetically labeled with myr::GFP (green) and immunostained with anti-Dpn (purple), Syp (cyan) and Imp (red). Note: the CNS was mounted laterally to visualize on the same section old vs young MNs (**see STAR method)**. **(L)** Bottom: schematic of the cell body of the 29 LinA/15 MNs. Top: schematic of the expression of Imp and Syp RBPs and of the expression of Chinmo, Mamo, Jim, Broad and Nvy proteins (horizontal bar, from **Fig.1**) and mRNA.

These results suggested that the post-transcriptional regulation plays a role in shaping the combinatorial code of TFs in iMNs.

### Opposite spatial gradients of Imp and Syp control the specificity of axonal targeting

To investigate post-transcriptional regulation further, we analyzed the expression dynamics and function on muscle innervation of the two known RNA binding proteins (RBPs), Imp and Syp two major players in the temporal specification of VNC and central brain lineages in Drosophila (Liu et al., 2015; Syed et al., 2017). In Lin A/15 NB and as has been described for other NBs in the CNS, both Imp and Syp proteins follow opposite temporal expression gradients (Wenyue *et al., unpublished*).

In iMNs, Imp and Syp proteins have opposite spatial gradients that are linked to their temporal expression in the NB: ventral (young) iMNs express a high level of Syp and a low level of Imp, whereas dorsal (older) iMNs express a high level of Imp and do not express Syp (**Fig. 4K**).By analyzing the coexpression of Imp and Syp in iMNs with Jim TFs **(Fig. 4J1-J4),** we concluded that Imp is highly expressed in iMNs 1 to 22 and weakly expressed in MN 23 to 29, whereas Syp is highly expressed in iMNs 23 to 29 and not detected in younger iMNs **(Fig. 4L)**. We hypothesized that their opposite expression patterns in iMNs could play a key role in shaping the axon-muscle connectome by establishing the TF code in iMNs.

We next examined the function of Imp and Syp on shaping the correct connectome between Lin A/15 axons and leg muscles **(Fig. 5)**. In *imp-/-* MARCM clones, the gene coding an inhibitor of apoptosis in the insect, the baculovirus P35 (Hay et al., 1994), was included in the genetic background to prevent MN death by apoptosis (Wenyue *et al., unpublished*) **(Fig. 5A1-A3, D1-D3)**. In this genetic background, the distal region of the tibia, which, in WT Lin A/15 is innervated by MNs expressing a low level of Imp during development, was not affected. By contrast, all the other regions were affected with variable penetrance (the distal region of the femur, tirm; the medial region of the tibia, talm and tadm; and the proximal region of the femur, tidm; **Fig. 5A1-A3, D1-D3 and Fig. S5)**. In *syp-/-* MARCM clones, a phenotype opposite to that of imp -/- was observed. The distal region of the tibia was not innervated by Lin A/15 axons (tarm1/2 and tadm), whereas other leg regions were innervated. Interestingly, innervation was greater than normal in the distal region of the femur (tirm) suggesting that abolishing Syp function is sufficient to re-direct axonal targeting from distal tibia to distal femur **(Fig. 5B1-B3, E1-E3)**. The ectopic expression of Imp in all Lin A/15 cells including the Syp+ cells (which normally express a low level of Imp) induced a similar innervation phenotype to that observed in *syp-/-* MARCM clones **(Fig. 5C1-C3, F1-F3)**. The result highlights the capacity of Imp to inhibit iMNs targeting of the distal tibia and to promote the targeting of the distal femur. Only a minor innervation phenotype (the talm is not targeted) resulted from the overexpression of Syp in Lin A/15 **(Fig. S5)**, suggesting a dorsal prevalence of Imp over Syp. Although, Imp and Syp negatively cross-regulate each other in the brain and VNC NBs (Liu et al., 2015; Syed et al., 2017), Imp/Syp mutual inhibition was not observed in our genetic manipulations in late L3 VNC, revealing that both RBPs act independently on specifying axonal targeting **(Fig. S6)**.

**Figure 5.**
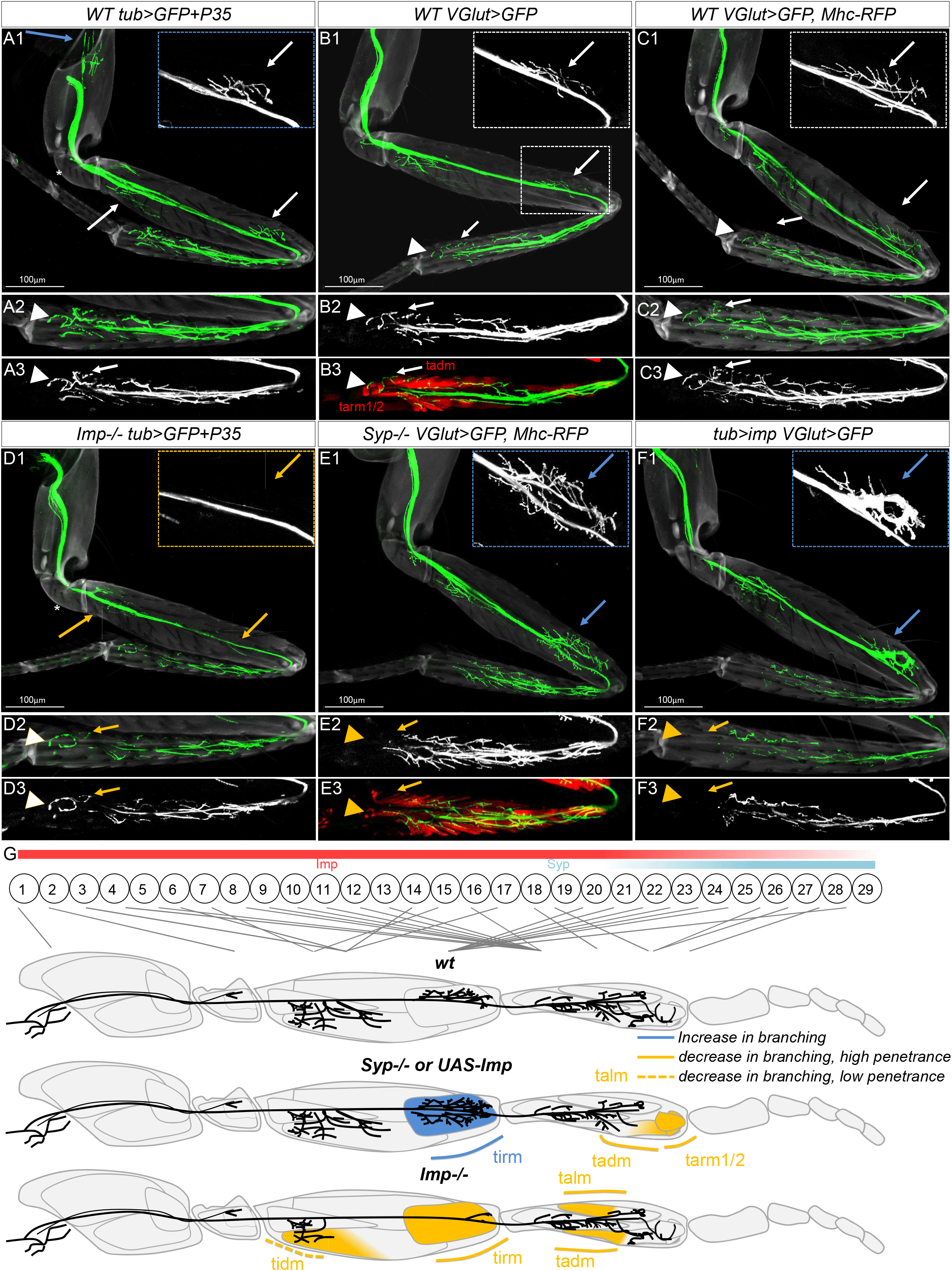
Opposite expression of Imp and Syp control the specificity of axonal targeting. **(A1-F3)** Axonal targeting phenotypes of *WT tub>P35* **(A1-A3)**, *Imp-/-, tub-P35* **(D1-D3)**, *WT* **(B1-B3, C1-C2)**, *Syp-/-* **(E1-E2)**, and *tub>imp* (**F1-F2**) Lin A/15 MARCM clones. The cuticle is in light grey and the axons are in green; muscles were labeled with *Mhc-RFP* (red) in **(B1-B3)** and **(E1-E3).** Insets show leg regions most affected. Arrowheads and arrows point normal targeting (white), the absence of/or reduced targeting (orange), and extra-targeting (blue). Note 1: asterisks indicate an absence of trochanter targeting in *Imp-/-, tub-P35* **(D1-D3)** that were not considered as part of the phenotypes because the *WT tub-P35* control showed similar defects **(A1-A3).** Note2: in *Syp-/-* **(E1-E3)** and *tub>imp* (**E1-E3**) Lin A/15 MARCM clones, the lack of muscle innervation in the distal tibia was not due to a decrease of MN number because more MNs were produced in MARCM clones **(see Fig S5)**, as a consequence of both the extended life of the NB and an inhibition of apoptosis in iMNs (Wenyue *et al.)*. Note 2: In *WT tub>P35* MARCM clones, the generated supernumerary neurons targeted a new region in the coxa (N=8/9). **(D1-D3)** N=9/12 (<u>N</u>umber of legs analyzed) for targeting defects at tarm1/2, N=6/12 for targeting defect at tadm in *Syp-/-* Lin A/15. **(E1-E3)** N=9/11 (<u>N</u>umber of legs analyzed) for targeting defects of the distal region of the tibia in Lin A/15 overexpressing Imp. **(F1-F3)** N=6/6 (<u>N</u>umber of legs analyzed) for targeting defects in the distal femur and N=2/6 for targeting defects in proximal region of the femur (tidm) region and medial region of the tibia (talm and tadm) in *Imp-/- tub>GFP+P35* **( See also Fig. S5 for *Imp-/- phenotypes*)**

In summary, our results show that Syp inhibits iMN targeting of the distal femur and promotes iMN targeting of distal tibia, whereas Imp has the opposite effect. Imp is required in early-born MNs for the correct targeting of the femur and proximal tibia. These results suggest that the opposite expression gradients of Imp and Syp in iMNs play a major role in shaping the architecture of the axon-muscle connectome.

### Imp and Syp shape the LinA/15 TF code

The function of Imp/Syp in shaping the axon-muscle connectome raised the question of whether they acted through the post-transcriptional regulation of the TF codes. To address this question, we focused on those 6 TFs whose expression has been characterized at the level of the RNA and Protein: Mamo, Chinmo, Broad, Nvy, Jim and Oli **(Fig.2, 4)**, and analyzed changes in their expression patterns when the levels of Imp and Syp were modified.

We tested the epistatic relationships between Imp/Syp and the TF codes in *syp-/-* and *Imp* overexpressing Lin A/15 clones because both genetic backgrounds induced similar innervation phenotypes and were therefore anticipated to induce similar variations in TF expression in the case of epistatic interactions. Five distinct effects were observed on TF expression. (i) Broad was expressed in fewer iMNs in the absence of Syp, **(Fig. 6A, D, J, Q)** whereas it was unaffected by Imp overexpression **(Fig. 6A, G, M, Q)**. (ii) Mamo expression was unaffected by the absence of Syp, but it was extended to more ventral iMNs with Imp overexpression **(Fig. 6A, I, O, Q; Fig. 6C, F, L)**. (iii) The expression of Chinmo and Jim were extended to ventral iMNs with both the absence of Syp and the overexpression of Imp **(Fig. 6A, D, E, G, H, J, K, M, N, P, Q)**. (iv) Nvy was expressed in fewer iMNs with both the absence of Syp and the overexpression of Imp **(Fig. 6B, E, H K, N, Q)**, Interestingly, only the ventral cluster of Nvy+ iMNs were affected by Imp overexpression, and not the dorsal cluster of Broad-Nvy+ iMNs **(Fig. S6)**. (v) Oli expression was unaffected by both the absence of Syp and the overexpression of Imp **(Fig. S6)**.

**Figure 6.**
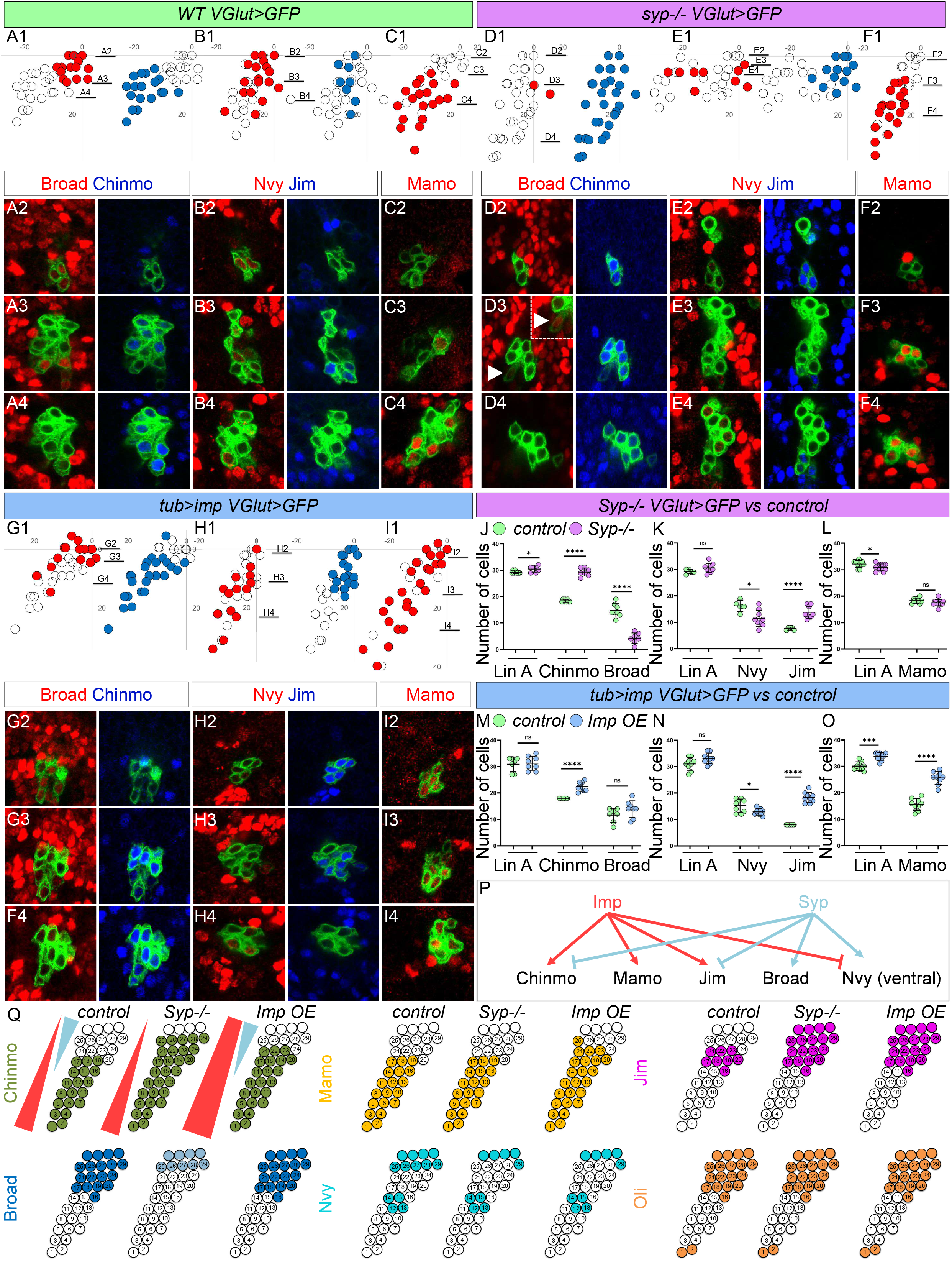
Opposite expression of Imp and Syp shapes the TF codes. **(A1-I4)** *WT* **(A1-C4)**, *Syp-/-* **(D1-E4)**, *tub>imp* (**G1-I4**) MARCM clones expressing mCD8::GFP (green) under the control *VGlut-LexA:GAD* and immunolabeled with anti-Broad (red) and anti-Chinmo (blue) **(A1-A4, D1-D4, G1-G4)** or anti-Nvy (red) and anti-Jim (blue) **(B1-B4, E1-E4, H1-H4)** or anti- Mamo (red) **(C1-C4, F1-F4, I1-I4)**. **(A2-A4, B2-B4, C2-C4, D2-D4, E2-E4, F2-F4, G2-G4, H2-H4, I2-I4)** are confocal sections from ventral to dorsal. **(A1, B1, C1, D1, E1, F1, G1, H1, I1)** Plots of the relative position of each Lin A/15 cell from a lateral perspective; the Lin A/15 cells expressing a given TF are color-coded in red or blue according to the immunostaining in **(A2-A4, B2-B4, C2-C4, D2-D4, E2-E4, F2-F4, G2-G4, H2-H4, I2-I4). (See also Fig S6 for Oli expression**, which was not affected). **(J-O)** Graph of the number of *VGlut+* Lin A/15 MNs in *Syp-/- VGlut>GFP vs WT VGlut>GFP* **(J-L)** *and in tub>imp VGlut>GFP vs WT VGlut>GFP* expressing Chinmo **(J, M)**, Broad **(J, M)**, Nvy **(K, N)**, Jim **(K, N)** and Mamo **(L, O)**. Of note: **(See also Fig S6 for Oli expression**, which was not affected). Note: the number of Lin A/15 *Vglut+* cells sometimes varied between controls and other genetic backgrounds. This was due the difficulty of having L3-larvae staged matched, correctly. However, the variations in **(J, O)** could not explain the large variation of the number of iMNs expressing Broad and Mamo. **(P)** Schematic of the epistasis between the Lin A/15 TFs and their upstream regulators Imp and Syp. **(Q)** Schematic of the iMN cell bodies of Lin A/15 from a lateral perspective in a third instar larva. The schematic shows the expression pattern of Chinmo, Mamo, Jim, Broad, Nvy and Oli in WT, *Syp-/-* and *Imp* overexpressing Lin A/15 MARCM clones. Note: only 33 iMNs are indicated because the read-out was *Vglut>GFP*, which was expressed in around 30 iMNs at that stage.

In summary, Imp and Syp had opposite effects on the expression of several TFs: Imp promoted the expression of Chinmo, Mamo and Jim proteins and inhibited the expression of Nvy whereas Syp inhibited the expression of Chinmo and Jim and inhibited the expression of Broad and Nvy **(Fig 6P, Q)**. These opposite effects of Imp/Syp on TF expression could explain why *syp-/-* and *Imp* overexpressing Lin A/15 induced similar innervation phenotypes.

### TF code is functional in building the axon-muscle connectome

We next analyzed the function of Chinmo, Broad, Nvy, Jim and Oli in shaping the axon-muscle connectome, using Lin A/15 MARCM knock-out or knock-down approaches (the attempts to analyze *mamo* were unsuccessful; **Fig. 7)**.

**Figure 7.**
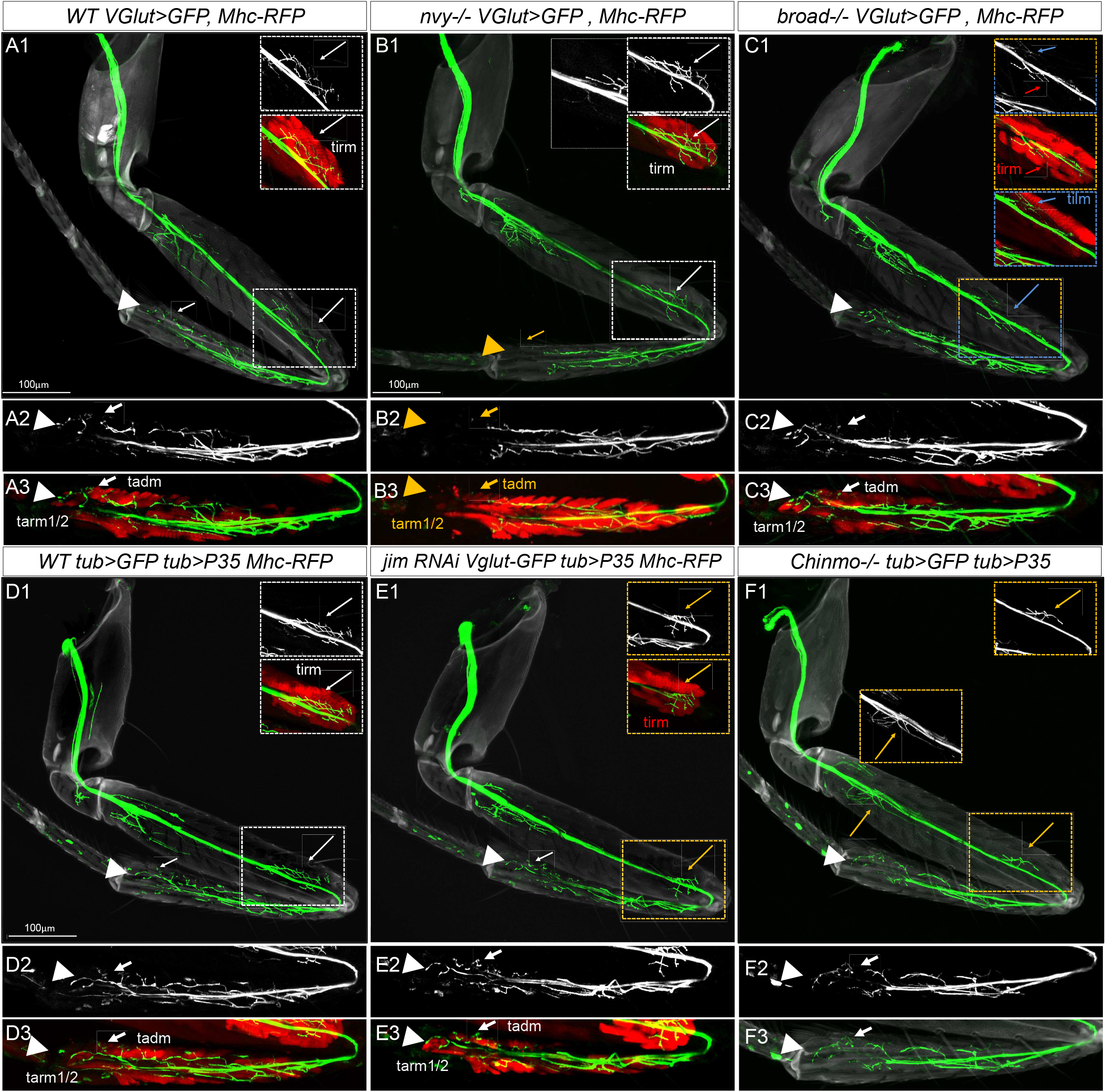
The TF codes shapes the axonal-muscle connectome. **(A1-F3)** Axon targeting phenotypes of *WT* **(A1-A3)**, *nvy/-* **(B1-B3)**, *broad-/-* (**E1-E2**), *WT, tub>P35* (**D1-D3)** *jim RNAi-/-, tub-P35* **(E1-E3)** and *chinmo-/-, tub-P35* **(F1-F3)** Lin A/15 MARCM clones expressing mCD8::GFP under the control of *VGlut* **(A1-C3, E1-E3)** or *tub* **(D1-D3, F1-F3).** The cuticle is light grey and the axons green; muscles were labeled with *Mhc-RFP* (red) in **(A1-E3)**. Insets show leg regions most affected. Arrowheads and arrows point normal targeting (white), the absence of or reduced targeting (orange), and extra-targeting (blue). Note 1: all pictures are max projections of confocal sections at the exception of the inset of the tirm and tilm that are partial max projections to highlight the muscle of interest. **(B1-B3)** the tarm1/2 in distal tibia (N=8/11), as well as the distal region of the tadm (N=2/11), were not innervated, and these phenotypes were often associated with a mistargeting of trlm in the coxa (N=7/11).

Nvy is expressed in 2 different MN clusters. One of the clusters correspond to the last-born MNs that mostly innervate the distal region of the tibia **(Fig. 1O)**. In *nvy-/-* clones, the distal region of the tibia is not innervated (tarm1/2 and distal region of the tadm) **(Fig. 7A1-A3, B1-B3)**. Broad is expressed in MNs targeting the distal femur and distal tibia (tadm, tarm1/2 the tirm) **(Fig. 2)**. In *broad -/-* clones, the tirm in the distal femur is less innervated, and is associated with a mistargeting of the tilm in the femur (N=9/22) (**7A1-A3, C1-C3)**. In *jim* knock-down clones, the tirm in the distal femur is less innervated by the 4 iMNs that normally express Jim and target the distal femur (N=13/17) (**7D1-D3, E1-E3)**. In *chinmo -/-* clones, the proximal (N=6/28) and distal (N=27/28) region of the femur were less innervated (**7D1-D3, F1-E3)**. Although the MNs targeting the proximal region of the femur normally express Chinmo, the MNs targeting do not, suggesting a non-autonomous function of Chinmo in these MNs **(see Discussion)**. The MNs expressing Oli TF are all affected with a variable penetrance in *oli-/-* Lin A/15 MARCM clones (**Fig S7**).

In summary, our results confirmed that the expression of Chinmo, Broad, Nvy, Jim and Oli in Lin A/15 iMNs specified axonal targeting. The epistatic relationships of 5 TFs with the two RPBs Imp and Syp imply that the two RBPs shape the architecture of the axon-muscle connectome through TFs.

### Post-transcriptional regulation of TFs in post-mitotic MNs by Imp shape the axon-muscle connectome

To establish the correct TF codes in iMNs, Imp and Syp could function in either the NB and/or iMNs. To discriminate between these two possibilities, we induced Imp expression in iMNs without affecting its temporal expression in the NB or GMC **(Fig. 8)**, using the *VGlut-gal4* driver.

**Figure 8.**
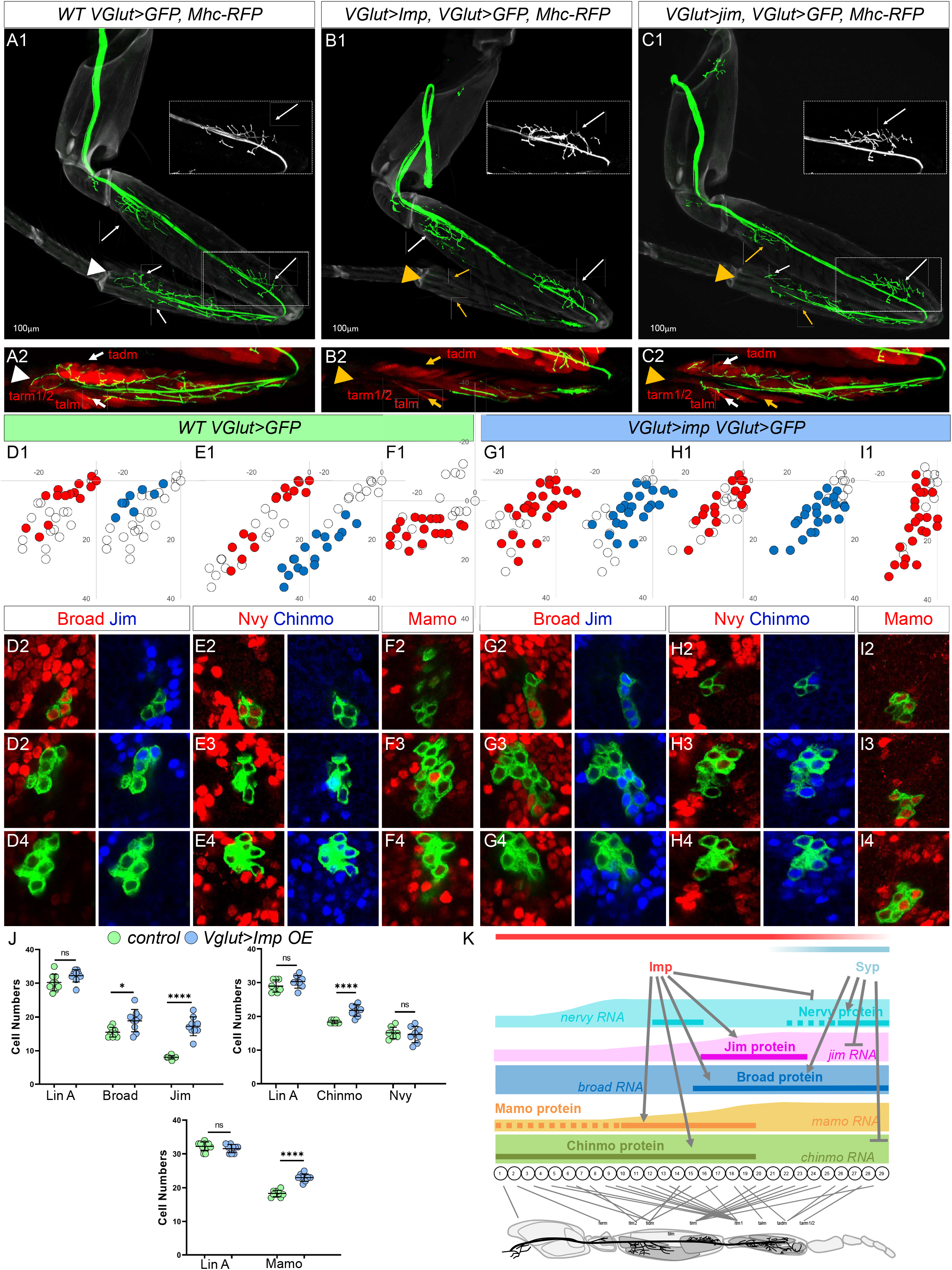
Post-transcriptional regulations of TFs in immature MNs by Imp shape the axon-muscle connectome. **(A1-C2)** Axon targeting phenotypes of *WT* **(A1-A2)**, *VGlut>Imp* **(B1-B2)** and *VGlut>Jim* (**C1-C3**), MARCM clones expressing mCD8::GFP under the control of *VGlut-Gal4*. The cuticle is light grey and the axons green; muscles were labeled with *Mhc-RFP* (red). Arrowheads and arrows point normal targeting (white) and the absence of or reduced targeting (orange). **(B1-B2)** In *VGlut>Imp* MARCM clones the tarm1/2 (N=8/11), tadm (N=4/11) and the talm (N=6/11) are not or less innervated (<u>N</u>umber of legs analysed). **(C1-C2)** In *VGlut>Jim* the tarm1/2 (N=12/14), the talm (N=10/14) and the tadm (1/14) are less innervated (Number of legs analysed), the proximal region of the femur is also less innervated (N=8/14) while the distal region of the femur is not affected. **(D1-I4)** *WT* **(D1-F4)** and *VGlut>Imp* **(G1-I4)** MARCM clones expressing mCD8::GFP (green) under the control of *VGlut-Gal4* and immuno-labeled with anti-Broad (red) and anti-Jim (blue) **(D1-D4, G1-G4),** or anti-Nvy (red) and anti-Chinmo (blue) **(E1-E4, H1-H4)** or anti-Mamo (red) **(F1-F4, I1-I4)**. **(D2-D4, E2-E4, F2-F4, G2-G4, H2-H4, I2-I4)** are confocal sections from ventral to dorsal; the confocal sections are indicated in **(D1, E1, F1, G1, H1, I1)**. **(D1, E1, F1, G1, H1, I1)** Plots of the relative position of each Lin A/15 cell from a lateral perspective; the Lin A/15 cells expressing a given TF are color-coded in red or blue according to the immunostaining in **(D2-D4, E2-E4, F2-F4, G2-G4, H2-H4, I2-I4).** **(J)** Graph of the number of *VGlut+* Lin A/15 MNs in *WT VGlut>GFP vs Vglut>imp, VGlut>GFP* expressing Chinmo, Broad, Nvy, Jim, and Mamo **(L, O). Note:** The increase of the number of MNs expressing Broad was observed affected the level of Imp generated the tub driver, but was not statistically significant (**Fig. 8J vs Fig. 6**). **(K)** Schematic of a model explaining how the architecture of the axon-muscle connectome is shaped. Imp and Syp post-transcriptionally refined the expression in iMNs of several TFs that in turn shape the axon-muscle connectome. Note: the doted and continuous lines indicate low and high expression levels of the TF proteins, respectively.

In this genetic background, muscle innervation phenotypes were similar to those generated by Imp overexpression in all Lin A/15 cell types, including the NB (**Fig. 8A-B** vs **Fig. 6**). In particular, the distal region of the tibia (tarm1/2, tadm and talm) was not or was less innervated than normal. Moreover, in this genetic background, similar epistatic relationships were observed between Imp and the TF codes to those observed by Imp overexpression in all Lin A/15 cell types (**Fig. 8D1-j** vs **Fig. 6**). In *VGlut>Imp* MARCM clones, the number of iMNs expressing Chinmo, Mamo, Jim and Broad was higher (**Fig. 8J**)., Although the number of iMNs expressing Nvy was not statistically lower in *VGlut>Imp* MARCM clones (**Fig. 8J**), it was in *tub>Imp* MARCM clones (**Fig. 6**). Interestingly, the overexpression of Jim only in iMNs, including last born MNs, induces a similar phenotype at the one induce in *VGlut>Imp* MARCM clones, and characterized by decreased innervation of the distal tibia (tarm1/2, tadm and talm) **(Fig. 8C)**.

These results show that the level of Imp shapes the axon muscle-connectome through the regulation of TFs in iMNs. Together with the above results, these observations allow us to propose a model where Imp and Syp shape the axon muscle connectome through the post-transcriptional regulation of TFs in iMNs **(Fig 8J, see Discussion)**.

## DISCUSSION

Several studies have revealed the functions of TF codes in specifying neuronal identities. However, the mechanisms that shape the expression of distinct TF codes at the single cell level remains largely unknown. In parallel to the transcriptional regulation of neuronal cell fate, recent pioneering work has revealed that post-transcriptional regulation plays a key role in the generation of different neuronal identities (Yang et al., 2014; Zahr et al., 2018). In our study, we have revealed an important relationship between transcriptional and post-transcriptional regulation for specifying neuronal cell fates. We have deciphered a molecular logic where the post-transcriptional regulation exerted by two known RBPs, Imp and Syp, contributes to progressively establishing the mTF codes in iMNs. In turn, the mTF codes specify individual morphological identities on the MNs during development.

### The axon-muscle connectome is shaped by a code of transcription factors expressed at the single cell level

Lin A/15 NB divides every 70 mins (Baek and Mann, 2009) and at each of the 29 divisions, it produces a GMC that generates an iMN with an unique developmental fate. Our immunostaining screening and computational analysis have demonstrated that the developmental fate of each Lin A/15 iMNs is associated with a TF code at the single cell level, and is established as a function of birth order before the iMN undergos morphological differentiation. The phenotypical analysis of muscle innervation when the TF expression is genetically modified **(Fig.6-8)** demonstrated that the TF codes specify the architecture of the axon-muscle connectome. These TFs have been termed mTFs because they determine the morphology of MNs (Enriquez et al., 2015). The precise characterization of the expression profiles of these TFs through development reveals the mTF codes are not established at each division but gradually shaped during development. Further, we show that post-transcriptional regulation by two RBPs, Imp and Syp, progressively shape the TF code.

This developmental logic for the regulation of MNs targeting contrasts with mushroom body (MB) neurons. A Lin A/15 NB produces few neurons with a high level of morphological diversity, whereas the MB NBs generate a high number of neurons with very little morphological diversity. The developmental process for MB neurons (which has been suggested to be more compatible with vertebrate neurogenesis) finds its molecular basis in the Chinmo TF gradient regulated by Imp and Syp. In Lin A/15, Imp and Syp are two major components of the post-transcriptional machinery shaping the TF code, demonstrating that the post-transcriptional regulation of TFs is also compatible with the neurogenesis of lineages generating a large amount of neuronal diversity in a short period of time. Here, we propose that the generation of 29 distinct fates requires the expression of mTF codes, in which expression patterns are controlled by two layers of regulators, transcriptional and post-transcriptional, that work in synergy to progressively sculpt the TF codes. However, the transcriptional upstream regulators inducing the expression of the mRNAs coding for the TFs were not identified. Theoretically, TFs expressed broadly in all iMNs are strong candidates to induce the expression of the mTF RNAs expressed ubiquitously. Interestingly some TF RNAs, such as *jim* or *mamo*, are expressed ubiquitously in a gradient. We propose that spatial selectors, such as the HOX TFs Antp or Ubx, are good candidates because their expression patterns follow a gradient in all Lin A/15 MNs, and are known to play a major roles in determining the identity of Lin A MNs (Baek et al., 2013). Other good candidates are the TFs found in our expression screen, such as RunxB, Islet or Jumu, that have a gradient expression pattern in Lin A/15 iMNs from high to low in ventral (young) and dorsal (old) iMNs, respectively. Finally we cannot exclude that tTFs could function in parallel with RBPs to regulate mTFs such as in larval MNs (Seroka et al., 2020).

### Imp/Syp function in parallel in iMNs to diversify their morphological identities

Imp and Syp play a central role in determining the fate of old versus young MNs. We propose that the morphological fate of each MN is in part, determined by an autonomous function of Imp/Syp in iMNs that shapes the mTF codes (**Fig. 8**). Both proteins have opposite and synergetic effects on the mTF RNAs. Imp promotes the translation of *chinmo, mamo, jim*, but inhibit the translation of *nvy*. By contrasts, Syp inhibits the translation of *chinmo* and *jim* and promots the translation of *nvy*. Interestingly both RBPs promote the translation of *broad*. These epistatic relations between Imp/Syp and the mTF codes explain the phenotypes of the axon-muscle connectome. In both *syp-/-* or Imp overexpressing *(UAS-Imp)* MARCM clones, Nvy is downregulated and Jim upregulated in the iMNs that target distal tibia and the innervation this region is reduced. The innervation in the distal tibia was also inhibited in *nvy-/-* and *jim RNAi* Lin A/15 MARCM clones.

Both RBPs have been described to cross inhibit in other CNS lineages (Liu et al., 2015; Syed et al., 2017; Yang et al., 2017a). However, in our genetic experiments, RBPs appeared not to cross inhibit, even though they acted in opposing directions to shape the axon-muscle connectome (**Fig. S6**). Our results suggested that mTFs are differentially sensitive to the relative concentration of the two RBPs. Recent studies have reveal that the interaction of the RBPs on RNAs coding for TF code is probably direct. For example, jim, *mamo* and *prospero* RNAs are known to immunoprecipitate with Imp (Samuels et al., 2020). Investigating further into the mechanism controlling the Imp/Syp-mTFs molecular interactions could be a good entry point to improve our understanding on how a distinctive TF code is refined by the two RBPs. Moreover, our results also show that not all mTFs are sensitive to Imp/Syp concentration. For example, Oli is not affected when the expression level of Imp/Syp is modified suggesting that other RBPs remain to be identified.

Finally, while our work emphases on the role of Imp/syp in iMNs, both RBPs might also be important for the temporal identity of the NB, for example through the regulation of Chinmo in the NB. This could explain the decrease in tirm innervation in *chinmo-/-* Lin A/15 clones that we interpret as a function of Chinmo in the NB (tirm is a muscle innervated by MNs not expressing Chinmo).

### Imp/Syp are multi-tasked proteins determining several facets of neuronal identity

In our study we have shown that Imp and Syp are important to determine one aspect of the MN identity, the morphology. However, both proteins also determine other aspects of MN fate. In our parallel study, we have demonstrated that both RBPs act autonomously in iMNs to determine the number of MNs produced by Lin A/15 (Wenyue *et al., unpublished*). Syp prevents the survival of iMNs whereas Imp promotes it. Hence Imp and Syp are multitasked proteins that determine the fate of iMNs by controlling neuronal survival and morphology.

Interestingly both proteins might also control molecular neuronal features such as the expression of neuronal transmitter. We were able to generate Lin A/15 *Imp-/-* MARCM clones with a pan-cellular *tub-Gal4* driver but not with *VGlut-gal4*, an enhancer trap transgene of the gene coding for the vesicular glutamate transporter that is expressed by all *Drosophila* MNs (Mahr and Aberle, 2006). These results suggest that Imp might be also involved in the terminal molecular identity of MNs that define them as glutamatergic neurons. The transcription factors controlling these molecular features are called terminal selectors (Allan et al., 2005; Eade et al., 2012; Hobert, 2011, 2016). In Lin A/15, all MNs are glutaminergic, and hence, terminal selectors should be expressed ubiquitously in all Lin A/15 MNs. Good candidates for terminal selectors could be genes found in our screen and expressed in all iMNs such as Lim 3 and Nkx6. Interestingly in the MB, the TF Mamo, which is under the positive control of Chinmo and Syp, control terminal neuronal features of MB neurons (Liu et al.) (of note: in Lin A/15 iMNs, Chinmo and Syp did not regulate Mamo expression).

### From stem cells to immature neurons

How a temporal gradient of RBPs in the Lin A/15 NB is translated into a spatial gradient in ventral (young) vs dorsal (old) immature iMNs remains unclear. It has been proposed that the temporal gradient of the RBPs in the NB, modulated by extrinsic factors, is bequeathed in the progeny such as maternal genes in the drosophila embryo (Liu et al., 2015). However, the expression patterns of *imp* and *syp* in Lin A/15 suggest that other mechanisms shape their expression gradients in iMNs. Intronic smFISHs against *imp* and *syp* reveal that *both* RBPs are actively transcribed in iMNs **(Fig S8)**. These results suggest that the temporal expression of RBPs in the NB is not passively inherited by iMNs but directly acquired during development by iMNs. We propose that iMNs, as is known for the NB, can directly integrate external cues such as the ecdysone hormone or activins (Rossi and Desplan, 2020)(Homem et al., 2014) to modutale Imp and Syp expression, and this in turn could affect the mTF code. This hypothesis implies that iMNs are plastic enough to integrate external cues. Interestingly, this plasticity is revealed by the changes to neuronal fate and the mTF code in Lin A/15 overexpressing Imp **(Fig. 8)**.

Finally, our results show that iMNs have the capacity to change their morphologies, thus revealing that iMNs are multipotent cells in term of their potential to enter into different morphological fate. For example, in both *syp-/-* or Imp overexpressing Lin A/15, MNs targeting distal tibia are morphologically re-specified to target the distal femur. Our smFISH results revealed the molecular basis of this plasticity. Clusters of MNs have the potential to acquire many morphologies because they share a similar TF transcriptome and a second layer of post-transcriptional regulation determine their final morphological fate by progressively refining the TF codes.

## EXPIRIMENTAL PROCEDURES

### Key Resources Table

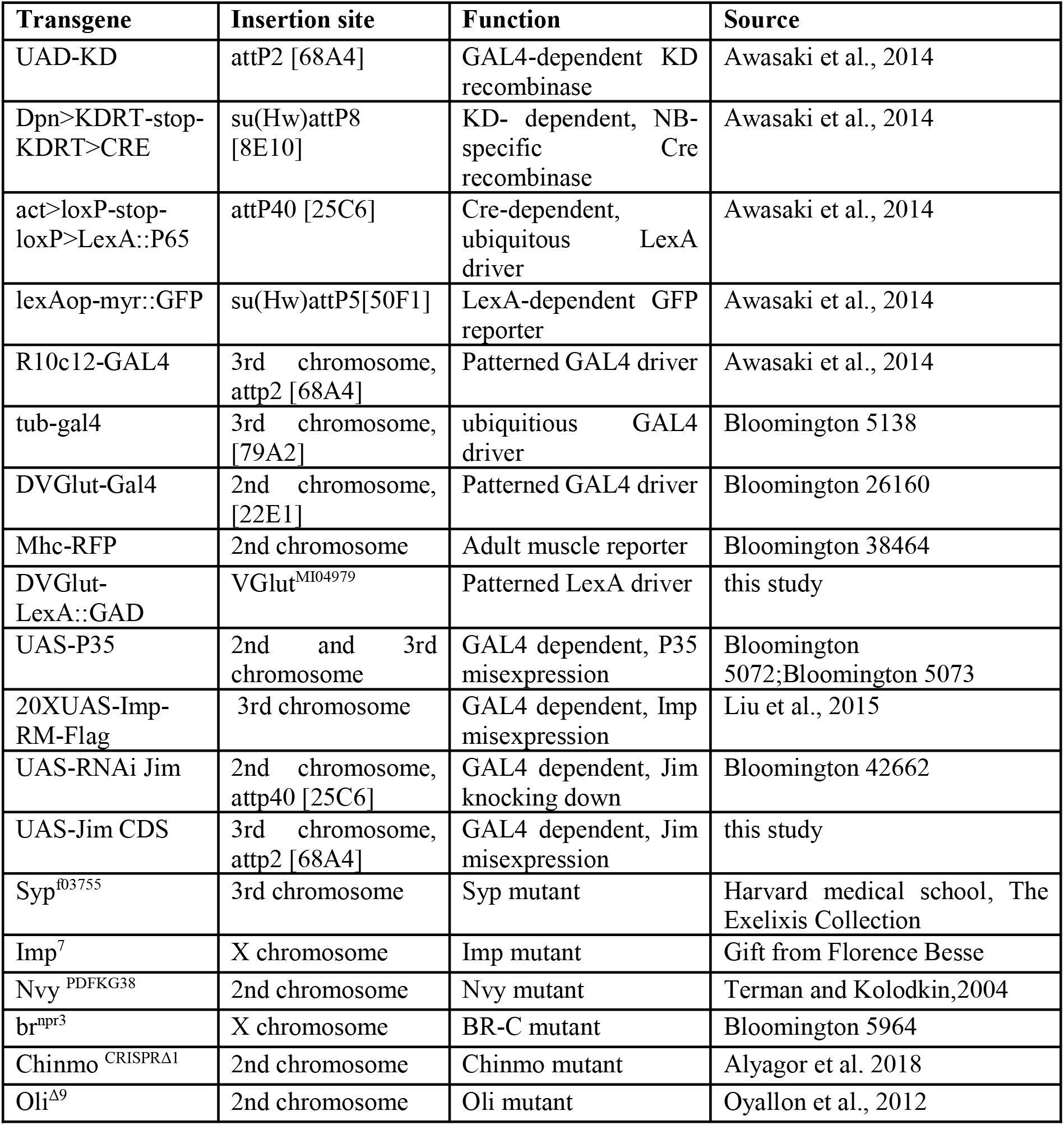

### Contact for Reagent and Resource Sharing

Further information and requests for resources and reagents should be directed to and will be fulfilled by the lead contact, jonathan enriquez.

### Experimental Model and Subject Details

All *in vivo* experiments were carried out using standard laboratory strains of *D. melanogaster*. See List of Transgenes and Genetic Crosses For each Figure for details. For stocks made in this study, the nucleotide sequences of the plasmids will be provided upon request.

#### Genetic Crosses for each Figure

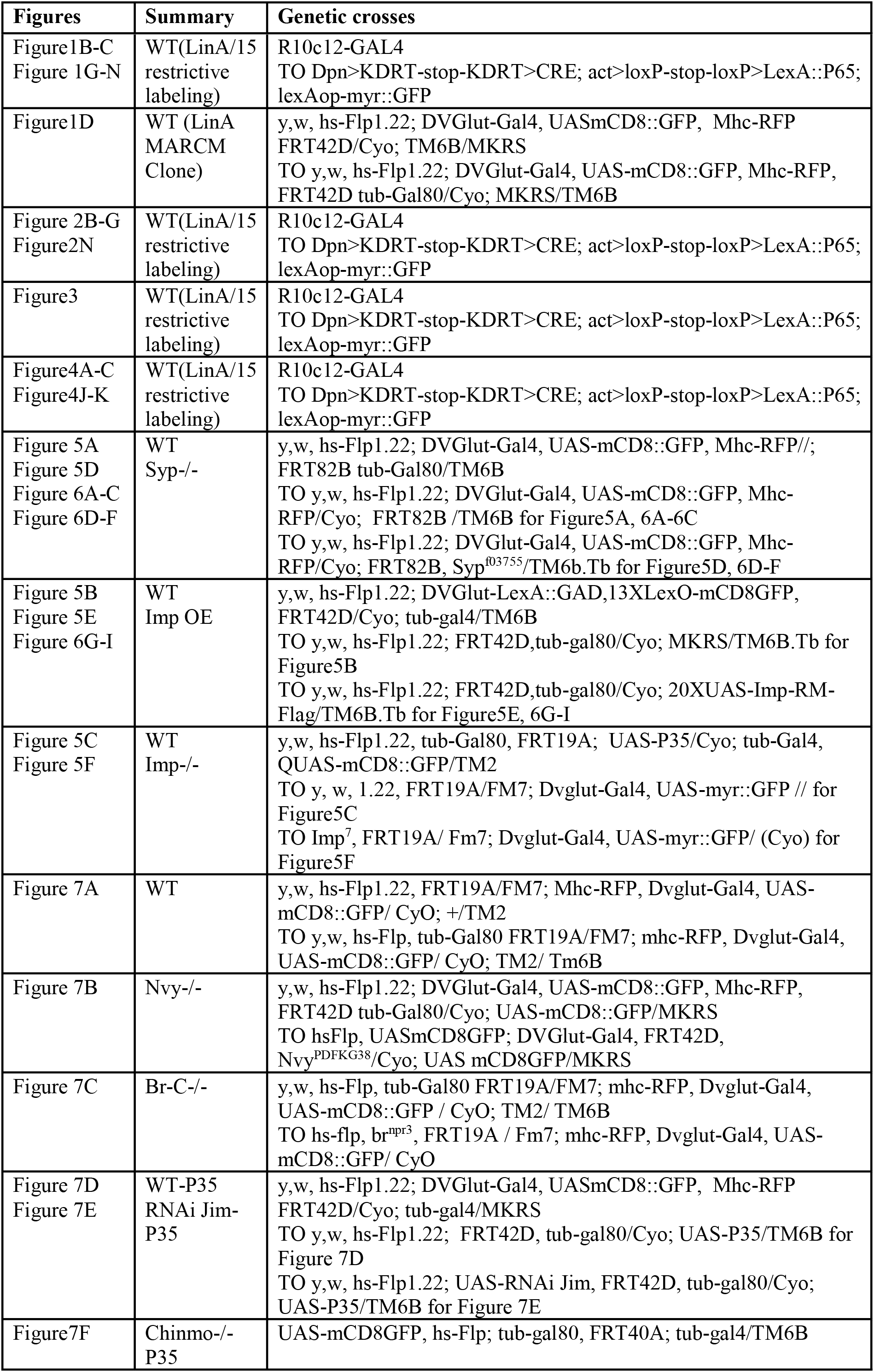
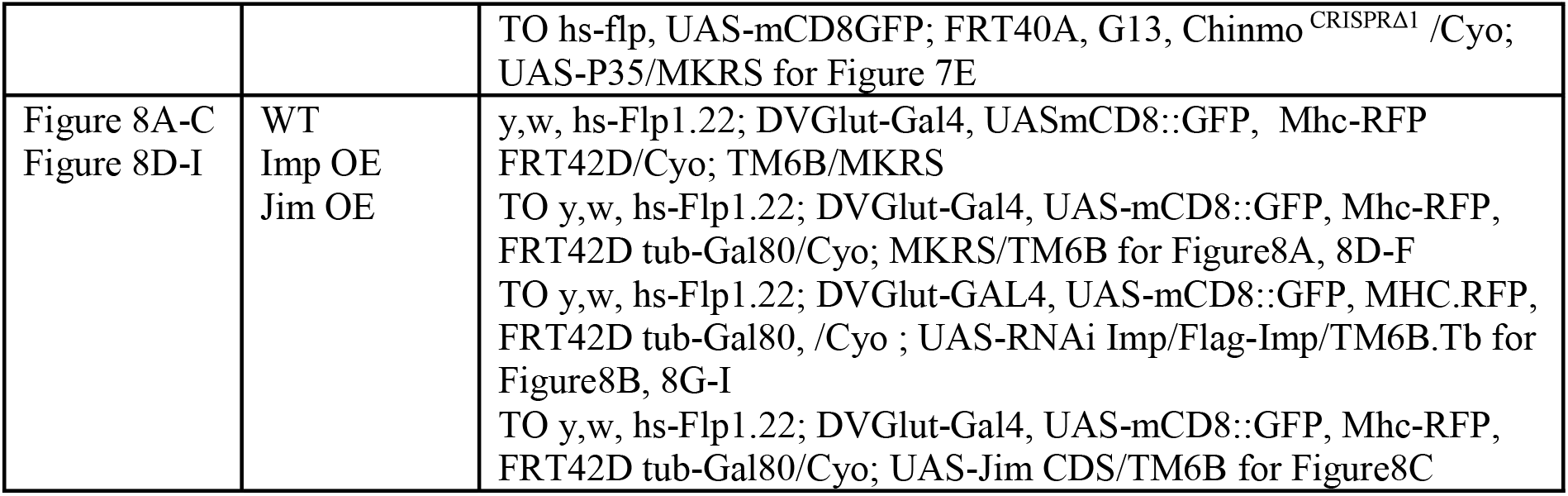

### Lineage tracing and MARCM Clonal Analysis

LinA/15 restrictive labeling is achieved by immortalizing Gal4 expression in Lin A/15 neuroblasts and its descendants (Awasaki et al. 2014; Lacin and Truman. 2016). Fly strains used to specifically label LinA/15 are listed in ***Genetic crosses for each figure***.

All fly strains used to introduce MARCM (Lee and Luo, 1999) clones labeling LinA/15 are listed in ***Genetic Crosses for each Figure***. To introduce MARCM clones, embryos were collected for 12hs in vials and incubated for 24hs at 25°C. First-instar larvae (0~12h ALH) were heat shocked at 37°C for 20 minutes to induce mosaic clones in L3 larvae and at 35°C for 15min to induce mosaic clones in adults. All flies were then raised at 25°C degrees at the exception of *Imp -/- tub>GFP+P35* flies being raised at 29°C after heat shock to boost the efficiency of P35 **(Fig.5F)**.

### Leg imaging

Legs were dissected, fixed and imaged as described in (Guan et al., 2018).

### Immunostaining of larval CNS

Inverted L3 larvae were fixed in 4% paraformaldehyde in PBS for 20 minutes at room temperature and blocked in the blocking buffer for one hour. L3 larval or pupal CNS were carefully dissected in PBS and then incubated with primary antibodies overnight (≥12h) and secondary antibodies in dark for one day (≥12h) at 4°C. Fresh PBST-BSA (PBS with 0.3% Triton X-100, 1% BSA) was used for the blocking, incubation, and washing steps: five times for 20 minutes at room temperature after fixation and after primary/secondary antibodies. Larval/pupal CNS were mounted onto glass slides using Vectashield antifade mounting medium (Vector Labs). Slides were either imaged immediately or stored at 4°C.

### Immunostaining of adult VNC

After removing the abdominal and head segments, the thorax of the flies were opened and fixed in 4% paraformaldehyde in PBS for 25 minutes at room temperature and blocked in the blocking buffer for 1 hour. After dissection, adult VNC were incubated with primary antibodies for 1 day and secondary antibodies in dark for 1 day at 4°C. Fresh PBST-BSA (PBS with 0.3% Triton X-100, 1% BSA) was used for the blocking, incubation, and washing steps: five times for 20 minutes at room temperature after fixation and after primary/secondary antibodies. VNC were mounted onto glass slides using Vectashield anti-fade mounting medium (Vector Labs). Slides were either imaged immediately or stored at 4°C.

### Primary and secondary antibodies

The initial set of antibodies were mostly generated by the modENCODE project or provided to us by C. Desplan (Li et al., 2013). About 250 antibodies were used and around 10 CNSs were analyzed for each antibody. Antibodies used for the 15 TFs are: rabbit anti-castor (1/150), mouse anti BR-C (1/25) (DSHB-25E9.D7), guinea-pig anti-Jim (1/100), mouse anti-prospero (1/100) (DSHB-MR1A), guinea-pig anti-Runt (1/100), rabbit anti-Nvy(1/1000), rabbit anti-RunXA (1/100), guinea-pig anti-FoxP (1/100), guinea-pig anti-Lov (1/100), rabbit anti-Zfh1(1/400) (gift from Jim Skeath), guinea-pig anti-Kr(1/300), Rat anti-Zfh2(1/250), rabbit anti-Oli (1/100), rabbit anti-Mamo (1/5000) (gift from Oren Schuldiner), guinea-pig anti-Chinmo (1/100).

Other primary antibodies used in this study include: mouse anti-Elav (1/100) (DSHB-9F8A9), rat anti-Elav (1/100) (DSHB-7E8A10), guinea-pig anti-Dpn (1/500) (gift from Jim Skeath, Skeath et al., 2017), rat anti-Imp (1/200) and rabbit anti-Syp (1/400) (gifts from Chris Doe).

The secondary antibodies used in this study include: goat anti-mouse Alexa 647 (Invitrogen-A32728), donkey anti-rat Alexa 647 (Jackson-712-605-153), goat anti-mouse Alexa 555 (Invitrogen-A32727), goat anti-rabbit Alexa 555 (Invitrogen-A32732), goat anti-Rat Alexa 555 (Abcam-ab150166), donkey anti-guinea-pig DyLight405 (Jackson-706-475-148).

### Image acquisition of immunostained VNC/CNS and adult legs

Multiple 0,5-μm-thick and 1-μm-thick sections in the z axis for larval/pupal CNS or adult VNC and adult legs respectively were imaged with a Leica TCS SP8 or a Zeiss LSM 780 confocal microscope. Objective used: 40x/1.3 oil objective for larval/pupal CNS or adult VNC; 20x glycerol objective for adult leg samples. Binary images for z stack images were generated using NIH ImageJ.

### 5-ethynyl-2’deoxyuridine (EdU) labelling

To mark MNs born at different time points (Figure 1K-M), L2 or L3 larvae (48, 72 and 96 hours after egg laying [AEL]) were transferred from standard fly food to fly food containing 250 mM EdU. Larvae were dissected at 120 hours AEL and dissected CNS were fixed in 4% paraformaldehyde in PBS for 20 minutes at room temperature, followed by a quick wash with PBST (PBS with 0.3% Triton X-100). Edu labeling was then detected using Clicl-iT EdU imaging kit (Invitrogen) according to manufacture’s instructions. An immunostaing was then performed as described in the **Immunostaining** section.

### Probe design and preparation for smiFISH

Primary probes against mRNA sequences of genes of interest (up to 48 probes per gene) were designed using the Biosearch Technologies stellaris RNA FISH probe designer tool (free with registration, https://biosearchtech.com). The probe sequences for each gene used in this study are shown in **(Supplemental File S1)**. The following sequence was added to the 5’ end of each 20 nuceotide (nt) probe: CCTCCTAAGTTTCGAGCTGGACTCAGTG, which is the reverse complement of the X flap sequence (CACTGAGTCCAGCTCGAAACTTAGGAGG) used in (Tsanov et al., 2016). The 5’-extened primary probe sets were synthesized by Integrated DNA Technologies (IDT), using 25 nmole synthesis scale, standard desalting, and at 100 μM in nuclease-free H2O. The X flap sequence, 5’ and 3’ end-labeled with Quasar 570, is synthesized by Biosearch Technologies. Fluorephore-labeled probe sets are prepared as described in (Tsanov et al., 2016), by hybridizing the fluorephore-labeled X Flap sequence and 5’-extented primary probe sets.

### smiFISH, sample preparation and hydrisation

Dissected larval CNS from third instar larvae were fixed in 4% paraformaldehyde (in PBS with 0.3% Triton X-100) for 20 minutes at room temperature. Samples were washed for 3 times of 15 minutes with PBST (PBS with 0.3% Tween-20) before a pre-hybridization wash in smiFISH wash buffer (10% deionised formamide (stored at −80°C) and 2x SSC in DEPC water) at 37°C for 30 minutes. Samples were then incubated with hybridized fluorephore-labeled probe sets diluted in smiFISH hybridization buffer (10% deionised formamide, 2x SSC and 5% dextran sulphate in DEPC water) at 37°C for 12 to 14 hours. The working concentration of each probe sets are: Jim (400 nM), Chinmo (800 nM), BR-C(1 μM), Nvy (2 μM), Mamo (2 μM) and Oli (400 nM). Samples were washed for 40 minutes at 37°C followed by three times of 15 minutes in smiFISH wash buffer at room temperature, and 10 minutes of washing in PBST (PBS with 0.3% Tween-20) before sample mounting. Protocol adapted from (Yang et al., 2017b).

### Image acquisition of larval VNC after smFISH

Images were acquired on a Leica TCS SP8 microscope using a 40x/1.3 oil objective. Image stacks were taken with the following settings: format 4400×4400 (223,56μm * 223,56μm (XY) and 44,95μm (Z)), speed 700 Hz, unidirectional, sequential line scanning, line averaging 8, pinhole 1 airy unit. Image stacks were acquired with a 350 nm z interval. Parameters for the three channels used in this study are: DAPI excitation 405 nm, laser 2%, detection 412-496nm; GFP excitation 488nm, laser 4.5%, detection 494-530nm; Quasar 570 excitation 548nm, laser power 5%, detection 555-605nm.

### SmFISH analysis

Each Lin A/15 cell was segmented in 3D in ImageJ/Fiji (Rueden et al., 2017; Schindelin et al., 2012; Schneider et al., 2012), using the LimeSeg plugin (Machado et al., 2019), on the GFP channel, with the following parameters: D_0~4-6, F pressure=0.01, Z_scale=6.8, Range in d0 units~4-6, Number of integration step=-1, real XY pixel size=50. For subsequent analysis, each segmented cell was exported into a separate ply file which was then imported in Matlab as a point cloud (The Math Works, Inc.). The original stacks were imported in Matlab using the Bio-Formats toolbox (https://www.openmicroscopy.org/bio-formats/downloads/). These stacks were then cropped around each cell using the point clouds generated by individual cell segmentation with LimeSeg.

The mRNA spots were detected in 3D, in the mRNA channel of these cropped stacks, using the method described by (Raj et al., 2008). In short, the spots were identified computationally by running a Matlab image processing script that runs the raw data through a filter (Laplacian of a Gaussian) designed to enhance spots of the correct size and shape while removing the slowly varying background. The filtered stacks are then thresholded to disregard remaining background noise. In order to choose an optimal threshold, all possible thresholds are computed. The thresholds were always chosen manually and close to the plateau. A ‘check File stack’ for each cell was generated in order to visualize the accuracy of the spot detection for a given threshold (**see supplemental experimental procedures S2**). In most of our samples, common thresholds were chosen for all the cells of a given Lin A/15. However, specific threshold were occasionally chosen for some cells. The parameters that gave the best visual detection of mRNA spots in our datasets-check files were generated were as follow: width of the Gaussian filter = 5; variance of the Gaussian filter = 0.25; threshold = 0.11 for *jim*, 0.08 for *chinmo, 0.15* for *mamo*, 0.09 for *broad, 0.29 for nvy* and 0.15 for Oli. The detected spots in each cell had a normal distribution of diameters with a mean of 0.25 +/- 0.05 micrometers and their maximum intensities displayed a unimodal distribution, arguing that the detected spots are mostly individual molecules. A custom Matlab routine transformed the point clouds corresponding to segmented cell volumes obtained from the LimeSeg plugin into an alpha shape (critical radius = 100) and then returned the number of mRNA contained in this cell volume using the Matlab function inShape.

### Quantification and statistical analysis

Graphs of the relative position of each LinA/15 cell were generated with Microsoft Excel. The spatial coordinates were assigned to each cell using the cell counter plug-in of NIH ImageJ software. The coordinates of each cell were normalized with Microsoft Excel in order to have the position of the Lin A NB at the origin of the plot graph. For samples where the NB was not labeled, the coordinates of each cell were then normalized to a cell located most anterior.

The plots of the number of TF-expressing cells (TFs are as indicated in each graph) were generated with Prism (GraphPad Software). All error bar represent standard deviation of a minimum of 7 Lin A/15, each dot represents a single Lin A/15 analyzed. Student’s t-test was performed to compare the difference in between indicated groups. Differences of P < 0.05 were considered significant. *0.01 < P < 0.05; **0.001 < P < 0.01; ***0.0001 < P < 0.001. **** P < 0.0001.

### Schematics

All schematic were done with Microsoft PowerPoint.

### Cloning

#### Pattb-20XUAS-hsp70-Jim-sv40

The full coding sequence of *jim* (2469bp) where the XhoI site in exon 1 was mutated to CTCCAG was ordered from Genewiz and cloned (XhoI/XbaI fragment) in the plasmid pJFRC165-20XUAS-IVS-R::PEST (https://www.addgene.org/32142/). The resulting *Pattb-20XUAS-hsp70-Jim-sv40* construct was inserted in position 86f on chromosome III by injection into embryos carrying *attP-86f* landing site.

### MIMIC swap to generate Mi{Trojan-VGlut-LexA::GAD.2}

#### Mi{Trojan-VGlut-LexA::GAD.2}

The plasmid *pBS-KS-attB2-SA(2)-T2A-LexA::GADfluw-Hsp70* was ordered from Addgene (https://www.addgene.org/78307/) and inserted in the MiMIC y[1] w[*]; Mi{y[+mDint2]=MIC}VGlut[MI04979] (BDSC_38078) line inserted into a coding intron in Phase 2 of the *VGlut* gene.

### Positive Cell Cluster Detection (PCCD) method (Figure 2, see supplemental note book 1)

The Positive Cell Cluster Detection (PCCD) method aims to link the expression of a given TF to the birth-order of a MN by using the correlation between the birth-order of MNs and their spatial organization. In our Lin A model, the Edu experiments **(Fig. 1)** reveal a good correlation between the birth order of MNs and their spatial distance from the NB in 3^rd^ instar larva: young born MNs are farer away from the NB compared to older MNs. The final goal of this method is to predict the TF code expressed in each immature MNs in a third instar larva.

The method followed a series of steps:

**Step 1:** From the imaging, assign spatial x, y, z coordinates and the expression (on/off) of a given TF to each Lin A cell (N>15, Number of Lin A immunostained for a given TF).
**Step 2:** Calculate the Euclidean distance between the NB and the x, y, z, coordinates of each iMN (relative distance).
**Step 3:** Order iMNs in each Lin A according to their distance to NB. This presents each Lin A as an ordered sequence of MNs (this defines the x axis position where cell #1 is defined as the furthest from the NB, i.e. the oldest MN on average). Then calculate the frequency of expression of all TFs as a function of their rank in each ordered Lin A sequence.
**Step 4:** Apply a filter (Savitzky-Golay) to smooth each distribution.
**Step 5:** Define the position in the sequence of the positive cell cluster(s) by using a peak detection method. Determine its length (average number of cells expressing a given TF in all the observed Lin A MN cluster expressing this TF). Then find the position of the positive cell cluster with this average length compatible with the smoothed TF distribution. The position and its length is represented by a horizontal line.
**Step 6:** Assemble all positive cell clusters for each TF on the same graph to reveal each TF combinatorial code. Convert the x’ axis to a birth order axis (1 to 29) since the distance between MN and the NB is tightly linked to their birth order. Define the coverage index at the border of all cell clusters.

More details about the method include:

**Savitzky-Golay filter (Step 4)**: The Savitzky-Golay algorithm (polynomial filter) was set with a window of size 11 and a polynomial order of 3 (see scipy.signal.savgol_filter function from the scipy python library).
**Peak detection method (Step 5):** Peaks were detected as local maxima in the normalized TF distributions. Local maxima were determined according to local conditions. They had a minimal height (h_min = 0.02), a minimal distance from other peaks (d_min = 15), and a minimal prominence (p_min = 0.1). The prominence of a peak measures how much a peak is emerging clearly locally in the signal. It is defined as the vertical distance between the peak and the altitude of the largest region it dominates. These values were found to yield best peak interpretations over the whole set of TF. We used the function scipy.signal.find_peaks of the scipy library.
**Positive cell clusters (Step 5):** For each TF, the average number p of positive cells was computed in each Lin A MN observed sequence. The procedure varied according to whether only one peak was detected or more than one (i.e. 2 in our data). Case of a single detected peak (e.g. Jim): The altitude of the peak at which the signal width below the peak is exactly p. The cluster of positive cells was assumed to correspond to all the cells expressing the TF. Case of two detected peaks (e.g. Oli). The sequence was split into the regions defined by by each peak. Then the average number of positive cells p1 and p2 are computed for each of the two regions. Then the method proceeds within each region and its average number of positive cells as in the case of a single detected peak. This determines both the estimated length and the position of the two positive cell clusters.
**Step 6**: The method leaves some uncertainty as to whether a cell located at the boundary of the cluster coverage should or not be considered as positive. If we consider that a cell at integer position x extends from x-0.5 to x+0.5, we can define a coverage index at the border (number from 0 to 1), revealing how much the positive cell cluster falls into the region around the cell (at −0.5 to + 0.5). For example, the length of the Jim positive cell cluster is 8.2 and its position is [14.8-23.1], which makes the two cells, cell 15 (14.8 belongs to [15-0.5,15+0.5] and cell 23 (23.1 belongs to [23-0.5,23+0.5], situated at the boundary uncertain of Jim expression. The coverage index is thus calculated for both cells, 0.7 (i.e: 15.5-14.8) for cell 15 and 0.6 (i.e: 23.1-22.5) for cell 23 (**Fig.2**). Since the average number of cells expressing Jim is 8.3, either cell, other than both, could be considered as positive and the method cannot distinguish between them. The coverage index at the border is indicated for cell cluster when inferior to 1.

## ACKNOWLEDGMENTS

We thank members of Alain Vincent, Filipe Pinto-Teixeira for comments on the manuscript. This work was funded by the Atip-Avenir program, FRM (#AJE20170537445) and AFM (#21999) to J.E, by the NIH: R01 NS070644 to R.S.M, and the GWIS-Nell Mondy Trust and Elizabeth Weisburger Fellowships to L.V. We acknowledge the contribution of SFR Biosciences (UAR3444/CNRS, US8/Inserm, ENS de Lyon, UCBL): Arthro-tool facility and of the IGFL facility platform.

## AUTHOR CONTRIBUTIONS

Conceptualization, J.E., W.G. and L.V.; Methodology, J.E., W.G and L.V.; Software, C.G. and J.E., Investigation, W.G., J.E., S.B., M.B., L.V., C.G., A.L., K.C., A.D., C.G., and S.U.; Writing – Original Draft, J.E.; Writing – Review & Editing, J.E., W.G., L.V., A.L., R.S.M., S.U., C.G., M.B.; Funding Acquisition, J.E.; Resources, J.E; Supervision, J.E and W.G.

## Notes

### Competing Interest Statement

The authors have declared no competing interest.

## REFERENCES

Allan, D.W., Park, D., St. Pierre, S.E., Taghert, P.H., and Thor, S. (2005). Regulators Acting in Combinatorial Codes Also Act Independently in Single Differentiating Neurons. Neuron 45, 689–700.

Alsiö, J.M., Tarchini, B., Cayouette, M., and Livesey, F.J. (2013). Ikaros promotes early-born neuronal fates in the cerebral cortex. Proc. Natl. Acad. Sci. 110, E716–E725.

Azevedo, A.W., Dickinson, E.S., Gurung, P., Venkatasubramanian, L., Mann, R.S., and Tuthill, J.C. (2020). A size principle for recruitment of Drosophila leg motor neurons. ELife 9, e56754.

Baek, M., and Mann, R.S. (2009). Lineage and birth date specify motor neuron targeting and dendritic architecture in adult Drosophila. J. Neurosci. Off. J. Soc. Neurosci. 29, 6904–6916.

Baek, M., Enriquez, J., and Mann, R.S. (2013). Dual role for Hox genes and Hox co-factors in conferring leg motoneuron survival and identity in Drosophila. Dev. Camb. Engl. 140, 2027–2038.

Bayraktar, O.A., and Doe, C.Q. (2013). Temporal patterning in intermediate progenitors increases neural diversity. Nature 498, 449–455.

Brierley, D.J., Blanc, E., Reddy, O.V., VijayRaghavan, K., and Williams, D.W. (2009). Dendritic Targeting in the Leg Neuropil of Drosophila: The Role of Midline Signalling Molecules in Generating a Myotopic Map. PLOS Biol. 7, e1000199.

Brierley, D.J., Rathore, K., VijayRaghavan, K., and Williams, D.W. (2012). Developmental origins and architecture of Drosophila leg motoneurons. J. Comp. Neurol. 520, 1629–1649.

Broihier, H.T., and Skeath, J.B. (2002). Drosophila Homeodomain Protein dHb9 Directs Neuronal Fate via Crossrepressive and Cell-Nonautonomous Mechanisms. Neuron 35, 39–50.

Broihier, H.T., Kuzin, A., Zhu, Y., Odenwald, W., and Skeath, J.B. (2004). Drosophila homeodomain protein Nkx6 coordinates motoneuron subtype identity and axonogenesis. Development 131, 5233–5242.

Certel, S.J., and Thor, S. (2004). Specification of Drosophila motoneuron identity by the combinatorial action of POU and LIM-HD factors. Development 131, 5429–5439.

Dasen, J.S., and Jessell, T.M. (2009). Chapter Six Hox Networks and the Origins of Motor Neuron Diversity. In Current Topics in Developmental Biology, (Academic Press), pp. 169–200.

Delile, J., Rayon, T., Melchionda, M., Edwards, A., Briscoe, J., Sagner, A., Klein, A., and Treutlein, B. (2019). Single cell transcriptomics reveals spatial and temporal dynamics of gene expression in the developing mouse spinal cord. Development 146.

Dillard, C., Narbonne-Reveau, K., Foppolo, S., Lanet, E., and Maurange, C. (2018). Two distinct mechanisms silence chinmo in Drosophila neuroblasts and neuroepithelial cells to limit their selfrenewal. Development 145, dev154534.

Doe, C.Q., and Skeath, J.B. (1996). Neurogenesis in the insect central nervous system. Curr. Opin. Neurobiol. 6, 18–24.

Eade, K.T., Fancher, H.A., Ridyard, M.S., and Allan, D.W. (2012). Developmental transcriptional networks are required to maintain neuronal subtype identity in the mature nervous system. PLoS Genet. 8, e1002501.

Elliott, J., Jolicoeur, C., Ramamurthy, V., and Cayouette, M. (2008). Ikaros confers early temporal competence to mouse retinal progenitor cells. Neuron 60, 26–39.

Enriquez, J., Venkatasubramanian, L., Baek, M., Peterson, M., Aghayeva, U., and Mann, R.S. (2015). Specification of Individual Adult Motor Neuron Morphologies by Combinatorial Transcription Factor Codes. Neuron 86, 955–970.

Enriquez, J., Rio, L.Q., Blazeski, R., Bellemin, S., Godement, P., Mason, C., and Mann, R.S. (2018). Differing Strategies Despite Shared Lineages of Motor Neurons and Glia to Achieve Robust Development of an Adult Neuropil in Drosophila. Neuron 97, 538–554.e5.

Fujioka, M., Lear, B.C., Landgraf, M., Yusibova, G.L., Zhou, J., Riley, K.M., Patel, N.H., and Jaynes, J.B. (2003). Even-skipped, acting as a repressor, regulates axonal projections in Drosophila. Development 130, 5385–5400.

Garces, A., and Thor, S. (2006). Specification of Drosophila aCC motoneuron identity by a genetic cascade involving even-skipped, grain and zfh1. Development 133, 1445–1455.

Guan, W., Venkatasubramanian, L., Baek, M., Mann, R.S., and Enriquez, J. (2018). Visualize Drosophila Leg Motor Neuron Axons Through the Adult Cuticle. J. Vis. Exp. JoVE.

Hay, B.A., Wolff, T., and Rubin, G.M. (1994). Expression of baculovirus P35 prevents cell death in Drosophila. Dev. Camb. Engl. 120, 2121–2129.

Hobert, O. (2011). Regulation of terminal differentiation programs in the nervous system. Annu. Rev. Cell Dev. Biol. 27, 681–696.

Hobert, O. (2016). Terminal Selectors of Neuronal Identity. Curr. Top. Dev. Biol. 116, 455–475.

Homem, C.C.F., Steinmann, V., Burkard, T.R., Jais, A., Esterbauer, H., and Knoblich, J.A. (2014). Ecdysone and Mediator Change Energy Metabolism to Terminate Proliferation in Drosophila Neural Stem Cells. Cell 158, 874–888.

Isshiki, T., Pearson, B., Holbrook, S., and Doe, C.Q. (2001). Drosophila neuroblasts sequentially express transcription factors which specify the temporal identity of their neuronal progeny. Cell 106, 511–521.

Jacob, J., Maurange, C., and Gould, A.P. (2008). Temporal control of neuronal diversity: common regulatory principles in insects and vertebrates? Dev. Camb. Engl. 135, 3481–3489.

Kohwi, M., and Doe, C.Q. (2013). Temporal Fate Specification and Neural Progenitor Competence During Development. Nat. Rev. Neurosci. 14, 823–838.

Konstantinides, N., Rossi, A.M., Escobar, A., Dudragne, L., Chen, Y.-C., Tran, T., Jaimes, A.M., Özel, M.N., Simon, F., Shao, Z., et al. (2021). A comprehensive series of temporal transcription factors in the fly visual system.

Landgraf, M., Roy, S., Prokop, A., VijayRaghavan, K., and Bate, M. (1999). even-skipped Determines the Dorsal Growth of Motor Axons in Drosophila. Neuron 22, 43–52.

Lee, T., and Luo, L. (1999). Mosaic Analysis with a Repressible Cell Marker for Studies of Gene Function in Neuronal Morphogenesis. Neuron 22, 451–461.

Li, X., Chen, Z., and Desplan, C. (2013a). Temporal patterning of neural progenitors in Drosophila. Curr. Top. Dev. Biol. 105, 69–96.

Li, X., Erclik, T., Bertet, C., Chen, Z., Voutev, R., Venkatesh, S., Morante, J., Celik, A., and Desplan, C. (2013b). Temporal patterning of Drosophila medulla neuroblasts controls neural fates. Nature 498, 456–462.

Liu, L.-Y., Long, X., Yang, C.-P., Miyares, R.L., Sugino, K., Singer, R.H., and Lee, T. Mamo decodes hierarchical temporal gradients into terminal neuronal fate. ELife 8.

Liu, Z., Yang, C.-P., Sugino, K., Fu, C.-C., Liu, L.-Y., Yao, X., Lee, L.P., and Lee, T. (2015). Opposing intrinsic temporal gradients guide neural stem cell production of varied neuronal fates. Science 350, 317–320.

Machado, S., Mercier, V., and Chiaruttini, N. (2019). LimeSeg: a coarse-grained lipid membrane simulation for 3D image segmentation. BMC Bioinformatics 20, 2.

Mahr, A., and Aberle, H. (2006). The expression pattern of the Drosophila vesicular glutamate transporter: a marker protein for motoneurons and glutamatergic centers in the brain. Gene Expr. Patterns GEP 6, 299–309.

Maniates-Selvin, J.T., Hildebrand, D.G.C., Graham, B.J., Kuan, A.T., Thomas, L.A., Nguyen, T., Buhmann, J., Azevedo, A.W., Shanny, B.L., Funke, J., et al. (2020). Reconstruction of motor control circuits in adult Drosophila using automated transmission electron microscopy. BioRxiv 2020.01.10.902478.

Mattar, P., Ericson, J., Blackshaw, S., and Cayouette, M. (2015). A Conserved Regulatory Logic Controls Temporal Identity in Mouse Neural Progenitors. Neuron 85, 497–504.

Maurange, C., Cheng, L., and Gould, A.P. (2008). Temporal Transcription Factors and Their Targets Schedule the End of Neural Proliferation in Drosophila. Cell 133, 891–902.

Okano, H., and Temple, S. (2009). Cell types to order: temporal specification of CNS stem cells. Curr. Opin. Neurobiol. 19, 112–119.

Oyallon, J., Apitz, H., Miguel-Aliaga, I., Timofeev, K., Ferreira, L., and Salecker, I. (2012). Regulation of locomotion and motoneuron trajectory selection and targeting by the Drosophila homolog of Olig family transcription factors. Dev. Biol. 369, 261–276.

Philippidou, P., and Dasen, J.S. (2013). Hox Genes: Choreographers in Neural Development, Architects of Circuit Organization. Neuron 80.

Prokop, A., and Technau, G.M. (1991). The origin of postembryonic neuroblasts in the ventral nerve cord of Drosophila melanogaster. Dev. Camb. Engl. 111, 79–88.

Raj, A., van den Bogaard, P., Rifkin, S.A., van Oudenaarden, A., and Tyagi, S. (2008). Imaging individual mRNA molecules using multiple singly labeled probes. Nat. Methods 5, 877–879.

Rossi, A.M., and Desplan, C. (2020). Extrinsic activin signaling cooperates with an intrinsic temporal program to increase mushroom body neuronal diversity. ELife 9, e58880.

Rueden, C.T., Schindelin, J., Hiner, M.C., DeZonia, B.E., Walter, A.E., Arena, E.T., and Eliceiri, K.W. (2017). ImageJ2: ImageJ for the next generation of scientific image data. BMC Bioinformatics 18, 529.

Sagner, A., and Briscoe, J. (2019). Establishing neuronal diversity in the spinal cord: a time and a place. Development 146.

Samuels, T.J., Järvelin, A.I., Ish-Horowicz, D., and Davis, I. (2020). Imp/IGF2BP levels modulate individual neural stem cell growth and division through myc mRNA stability. ELife 9, e51529.

Schindelin, J., Arganda-Carreras, I., Frise, E., Kaynig, V., Longair, M., Pietzsch, T., Preibisch, S., Rueden, C., Saalfeld, S., Schmid, B., et al. (2012). Fiji: an open-source platform for biological-image analysis. Nat. Methods 9, 676–682.

Schneider, C.A., Rasband, W.S., and Eliceiri, K.W. (2012). NIH Image to ImageJ: 25 years of image analysis. Nat. Methods 9, 671–675.

Seroka, A., Yazejian, R.M., Lai, S.-L., and Doe, C.Q. (2020). A novel temporal identity window generates alternating Eve+/Nkx6+ motor neuron subtypes in a single progenitor lineage. Neural Develop. 15, 1–14.

Skeath, J.B., Wilson, B.A., Romero, S.E., Snee, M.J., Zhu, Y., and Lacin, H. (2017). The extracellular metalloprotease AdamTS-A anchors neural lineages in place within and preserves the architecture of the central nervous system. Dev. Camb. Engl. 144, 3102–3113.

Soler, C., Daczewska, M., Ponte, J.P.D., Dastugue, B., and Jagla, K. (2004). Coordinated development of muscles and tendons of the Drosophila leg. Development 131, 6041–6051.

Syed, M.H., Mark, B., and Doe, C.Q. (2017). Steroid hormone induction of temporal gene expression in Drosophila brain neuroblasts generates neuronal and glial diversity. ELife 6, e26287.

Thor, S., and Thomas, J.B. (1997). The Drosophila islet Gene Governs Axon Pathfinding and Neurotransmitter Identity. Neuron 18, 397–409.

Thor, S., Andersson, S.G.E., Tomlinson, A., and Thomas, J.B. (1999). A LIM-homeodomain combinatorial code for motor-neuron pathway selection. Nature 397, 76–80.

Truman, J.W., and Bate, M. (1988). Spatial and temporal patterns of neurogenesis in the central nervous system of Drosophila melanogaster. Dev. Biol. 125, 145–157.

Tsanov, N., Samacoits, A., Chouaib, R., Traboulsi, A.-M., Gostan, T., Weber, C., Zimmer, C., Zibara, K., Walter, T., Peter, M., et al. (2016). smiFISH and FISH-quant – a flexible single RNA detection approach with super-resolution capability. Nucleic Acids Res. 44, e165–e165.

Venkatasubramanian, L., Guo, Z., Xu, S., Tan, L., Xiao, Q., Nagarkar-Jaiswal, S., and Mann, R.S. (2019). Stereotyped terminal axon branching of leg motor neurons mediated by IgSF proteins DIP-α and Dpr10. ELife 8, e42692.

Yang, C.-P., Samuels, T.J., Huang, Y., Yang, L., Ish-Horowicz, D., Davis, I., and Lee, T. (2017a). Imp and Syp RNA-binding proteins govern decommissioning of Drosophila neural stem cells. Development 144, 3454–3464.

Yang, G., Smibert, C.A., Kaplan, D.R., and Miller, F.D. (2014). An eIF4E1/4E-T complex determines the genesis of neurons from precursors by translationally repressing a proneurogenic transcription program. Neuron 84, 723–739.

Yang, L., Titlow, J., Ennis, D., Smith, C., Mitchell, J., Young, F.L., Waddell, S., Ish-Horowicz, D., and Davis, I. (2017b). Single molecule fluorescence in situ hybridisation for quantitating post-transcriptional regulation in Drosophila brains. Methods San Diego Calif 126, 166–176.

Zahr, S.K., Yang, G., Kazan, H., Borrett, M.J., Yuzwa, S.A., Voronova, A., Kaplan, D.R., and Miller, F.D. (2018). A Translational Repression Complex in Developing Mammalian Neural Stem Cells that Regulates Neuronal Specification. Neuron 97, 520–537.e6.

Zhu, S., Lin, S., Kao, C.-F., Awasaki, T., Chiang, A.-S., and Lee, T. (2006). Gradients of the Drosophila Chinmo BTB-Zinc Finger Protein Govern Neuronal Temporal Identity. Cell 127, 409–422.

