## supplemental information for "The post-transcriptional regulation of TFs in immature motoneurons shapes the axon-muscle connectome"

**Supplemental Figure and Figure Legends**

**Figure S1. Correlation between birth order and TF codes.**

(**A1, B1, C1, D1, E1, F1, G1, H1, I1, J1, K1, L1)** Plots of the relative position of the Lin A/15 cells from a lateral perspective in a late third instar lava. iMNs (blue), NB (cyan) , TF (red). The name the TF is indicated on the top of each plot.

**(A2-A3, B2-B3, C2-C3, D2-D3, E2-E3, F2-F3, G2-G3, H2-H3, I2-I3, J2-J3, K2-K3, L2-L3)** Confocal sections of the Lin A/15 in (**A1, B1, C1, D1, E1, F1, G1, H1, I1, J1, K1, L1**). Anti-Elav (blue), GFP (green), anti-Dpn (cyan), anti-TF (red). The name the anti-TF used is indicated on the top of each Plot. **(A2, B2, C2, D2, E2, F2, G2, H2, I2, J2, K2, L2)** are confocal sections trough the NB and **(A3, B3, C3, D3, E3, F3, G3, H3, I3, J3, K3, L3)** are confocal sections trough the TF+ cells.

**(O)** Schematic of the cell bodies of Lin A/15 in a third instar larva from a lateral perspective (not including the proliferative glia). The schematic shows the expression pattern of each TF (red) in **(A1-A3, B1-B3, C1-C3, D1-D3, E1-E3, F1-F3, G1-G3, H1-H3, I1-I3, J1-J3, K1-K3, L1-L3)** and is based on the PCCD method.

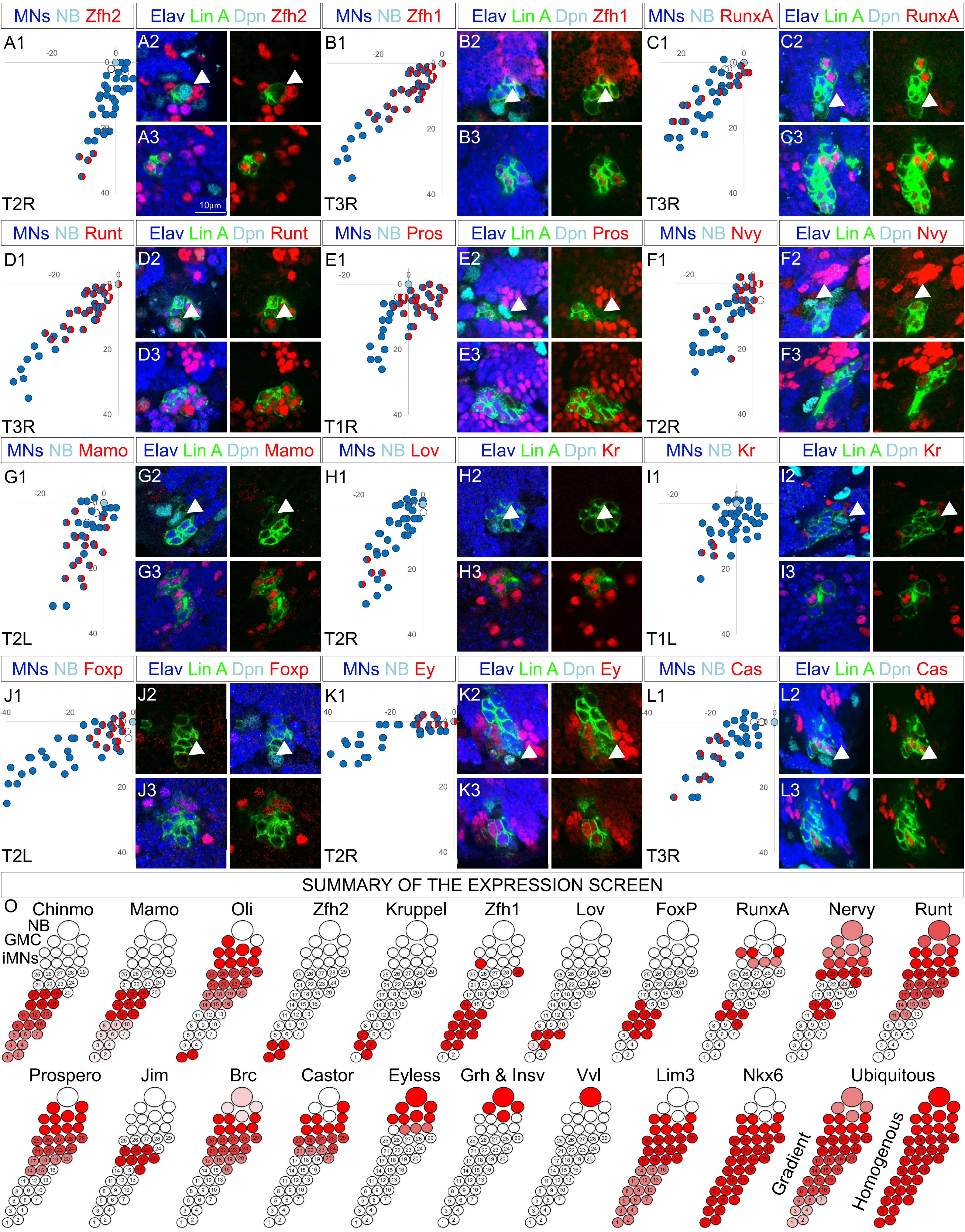


Gradient : RunXB, Antp, Islet, Jumu, Gt

homogenous: Psq; Seq; Ind; Adf1; E2F1; E2F2; Zelda; Sd; Snr1; Inv; ham; Hth; Tll; Grn; Ato, CG8108; CG12391; Dalao.

**Figure S1**

**Figure S2. Co-staining between TFs;**

**(A1-A2, B1-B2, C1-C2, D1-D3, E1-E2, F1-F2, G1-G2, H1-H2, I1-I3, J1-J2, K1-K3)** Confocal sections of Lin A/15 genetically labeled with myr:GFP and co-immunostained against pair of TFs in a third instar larva. The name of the two TF co-immunolabeled and the number of iMNs expressing both TF are written are written on the left of each set of confocal sections. **(A1, B1, C1, D1, E1, F1, G1, H1, I1, J1, K1, L1)** are ventral section compared to **(A2, B2, C2, D2, E2, F2, G2, H2, I2, J2, K2-3).**

**(A3, B3, C3, D3, E3, F3, G3, H3, I3, J3, K4)** Plot of the relative position of each Lin A/15 cell from a lateral perspective TF+ cells are in red and/or blue. The confocal sections in **(A1-A2, B1-B2, C1-C2, D1-D3, E1-E2, F1-F2, G1-G2, H1-H2, I1-I3, J1-J2, K1-K3)** are indicated by black lines.

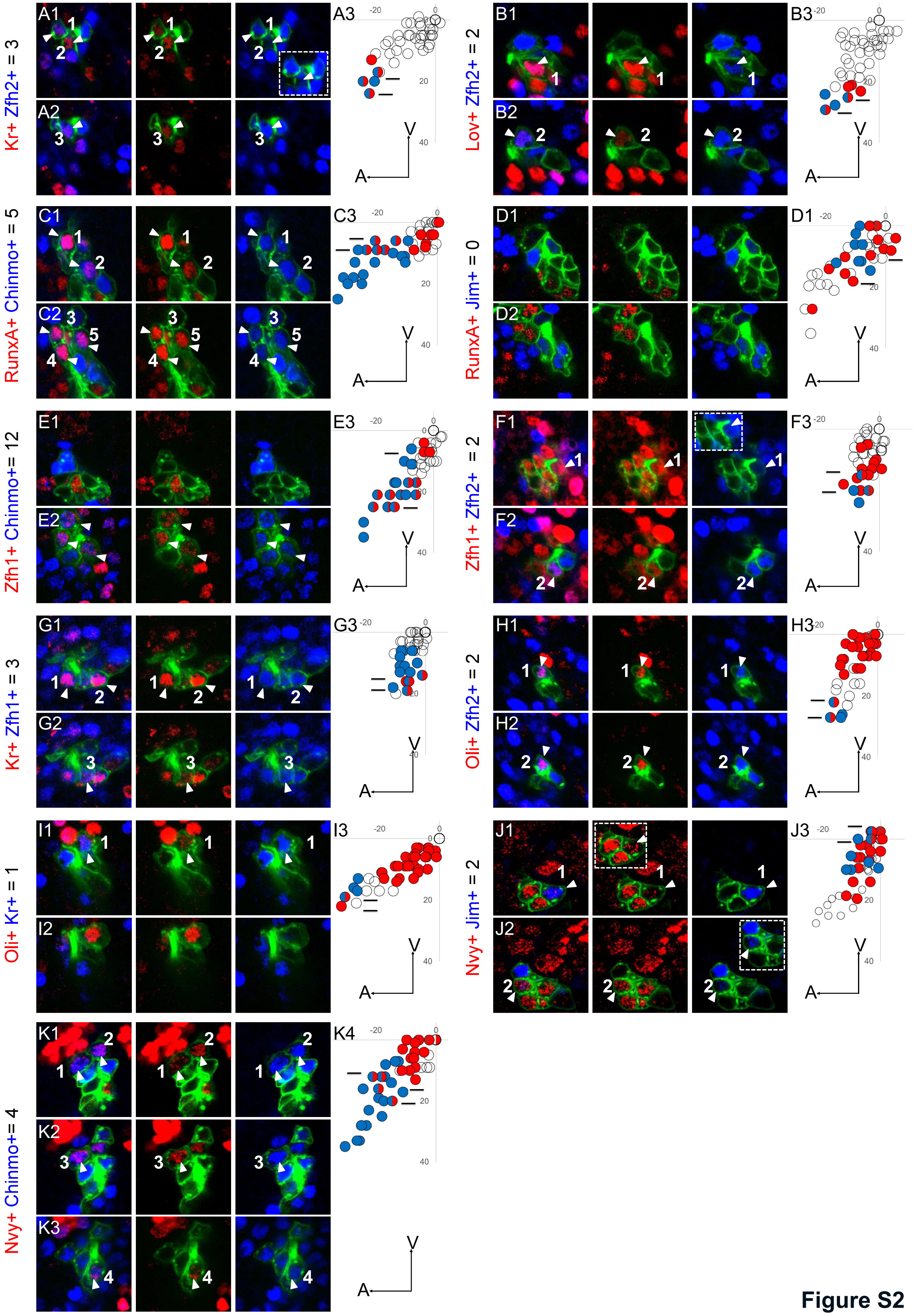

Kr+ Zfh2+ = 3
Lov+ Zfh2+ = 2
RunxA+ Chinmo+ = 5
RunxA+ Jim+ = 0
Zfh1+ Chinmo+= 12
Zfh1+ Zfh2+ = 2
Kr+ Zfh1+ = 3
Oli+ Zfh2+ = 2
Oli+ Kr+ = 1
Nvy+ Jim+ = 2
Nvy+ Chinmo+= 4


**Figure S2**

**Figure S3. The dynamic expression of the TF code**

**(A1, B1, C1, D1, E1, F1, G1, H1, I1, J1, K1, L1, M1, N1, O1, P1, Q1, R1, S1, T1, U1, V1)** Plots of the relative position of the Lin A/15 cells from a lateral perspective in early L3 (**A1, C1, E1, , G1, I1, K1, L1, M1, O1, Q1, S1, U1)** and mid L3 **(B1, D1, F1, H1, J1, L1, N1, P1, R1, T1, V1)**. iMNs (blue), NB (cyan), TF (red). The name the TF is indicated on the left of each plot.

**(A2-A3, B2-B3, C2-C3, D2-D3, E2-E3, F2-F3, G2-G3, H2-H3, I2-I3, J2-J3, K2-K3, L2-L3, M2-M3, N2-N3, O2-O3, P2-P3, Q2-Q3, R2-R3, S2-S3, T2-T3, U2-U3, V2-V3)** Confocal sections of the Lin A/15 in **(A1, B1, C1, D1, E1, F1, G1, H1, I1, J1, K1, L1, M1, N1, O1, P1, Q1, R1, S1, T1, U1, V1)**. Anti-Elav (blue), GFP (green), anti-Dpn (cyan), anti-TF (red). The name the anti-TF used is indicated on the left of each panel. (**A2, B2, C2, D2, E2, F2, G2, H2, I2, J2, K2, L2, M2, N2, O2, P2, Q2, R2, S2, T2, U2, V2)** are confocal sections trough the NB and (**A3, B3, C3, D3, E3, F3, G3, H3, I3, J3, K3, L3, M3, N3, O3, P3, Q3, R3, S3, T3, U3, V3)** are confocal sections trough  TF+ cells.

**(W)** Schematic of the cell bodies of Lin A/15 from a lateral perspective (not including the proliferative glia) at two developmental time points:  early-L3 and mid-L3. See figure S1 for late L3 time point. The schematic shows the expression pattern of each TF (red) in **(A1-A3, B1-B3, C1-C3, D1-D3, E1-E3, F1-F3, G1-G3, H1-H3, I1-I3, J1-J3, K1-K3, L1-L3, M1-M3, N1-N3, O1-O3, P1-P3, Q1-Q3, R1-R3, S1-S3, T1-T3, U1-U3, V1-V3)** and are categorized according their expression pattern.

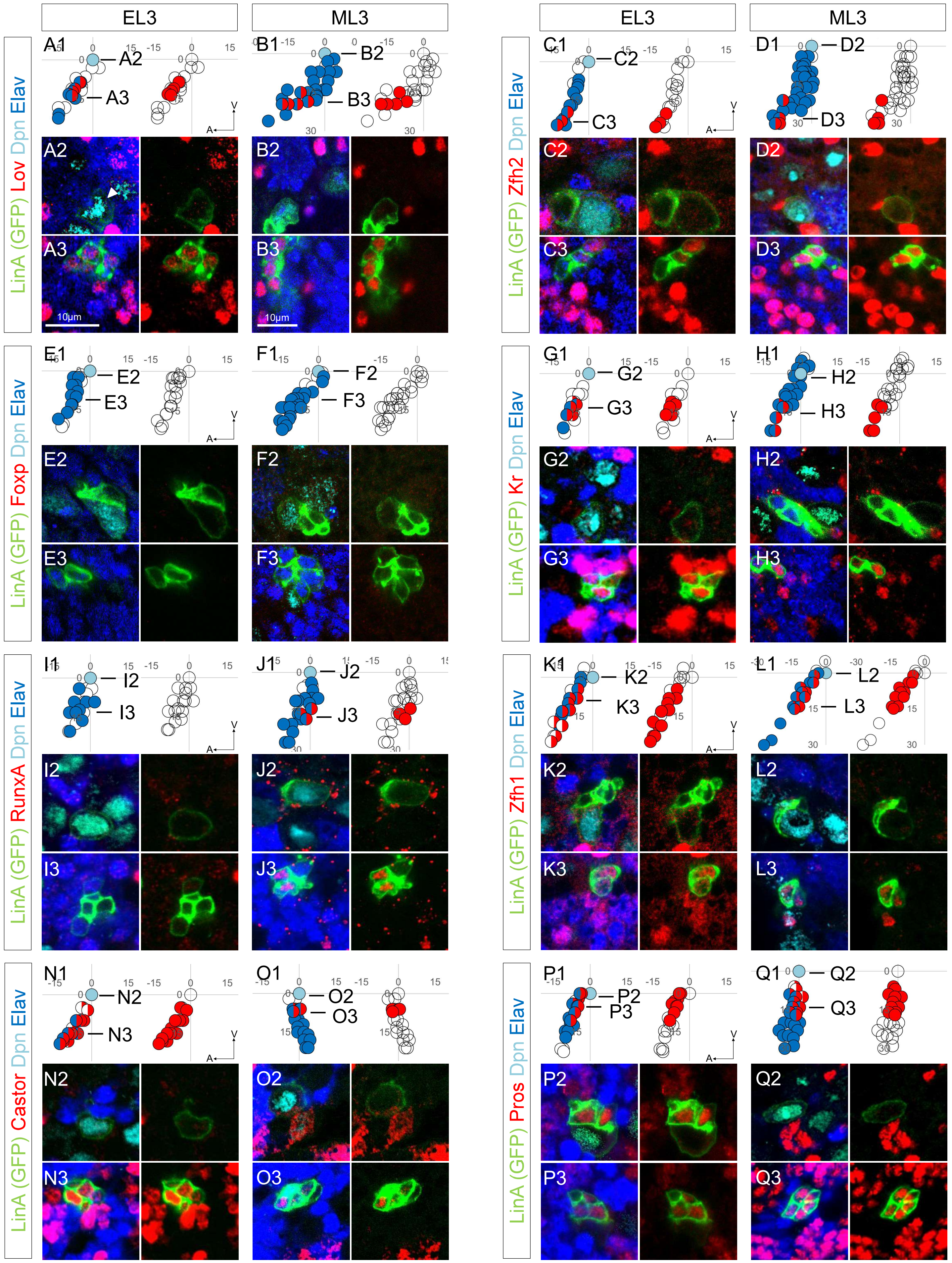
EL3
ML3
EL3
ML3
LinA (GFP) Lov Dpn Elav
LinA (GFP) Zfh2 Dpn Elav
LinA (GFP) Foxp Dpn Elav
LinA (GFP) Kr Dpn Elav
LinA (GFP) RunxA Dpn Elav
LinA (GFP) Zfh1 Dpn Elav
LinA (GFP) Castor Dpn Elav
LinA (GFP) Pros Dpn Elav
10μm
10μm

Figure S3

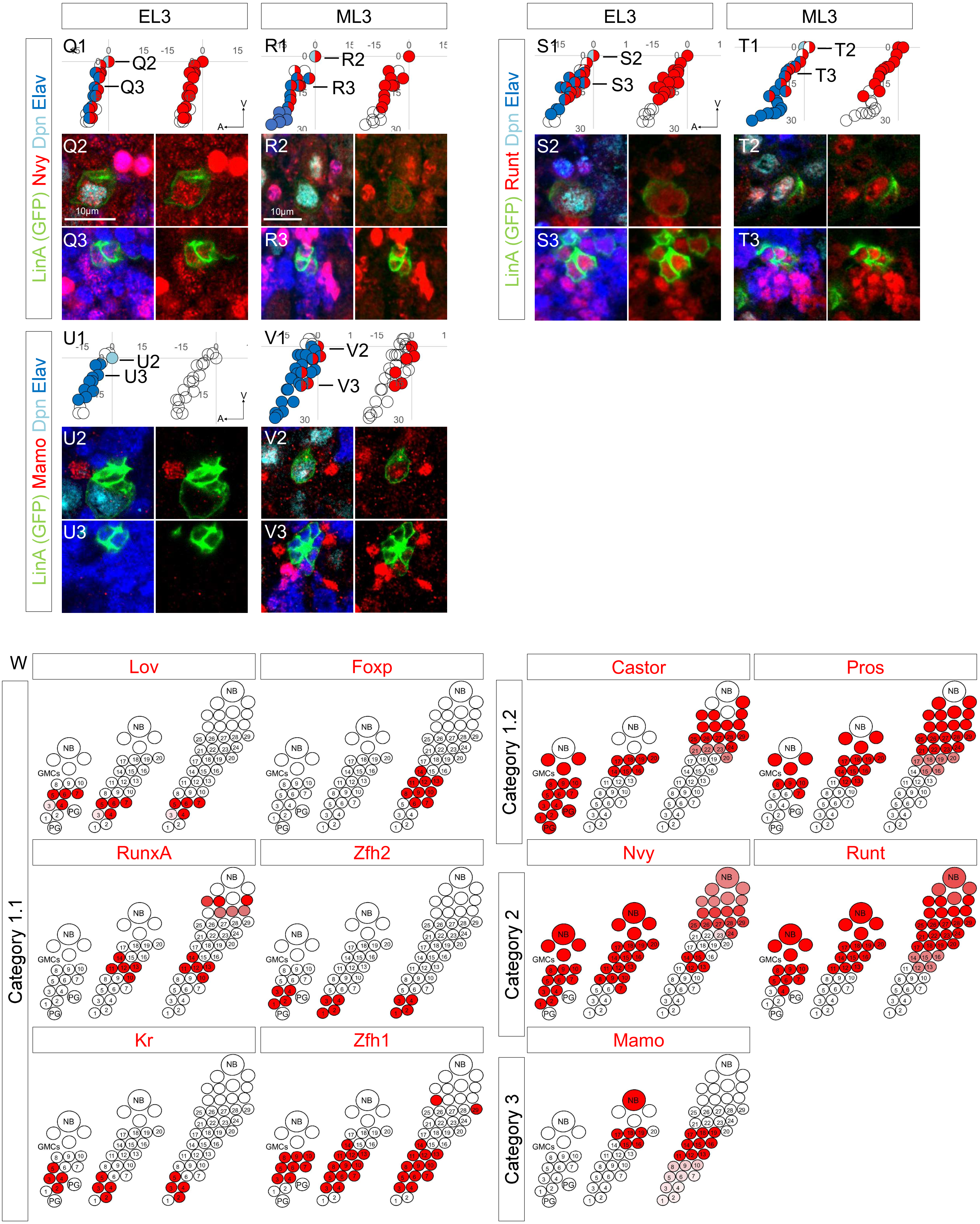
EL3
ML3
EL3
ML3
Q1
Q2
Q3
R1
R2
R3
S1
S2
S3
T1
T2
T3
LinA (GFP) Nvy Dpn Elav
LinA (GFP) Runt Dpn Elav
10µm
U1
U2
U3
V1
V2
V3
LinA (GFP) Mamo Dpn Elav
W
Lov
Foxp
Castor
Pros
RunxA
Zfh2
Nvy
Runt
Kr
Zfh1
Mamo
Category 1.1
Category 1.2
Category 2
Category 3
NB
GMCs
PG

Figure S3

**Figure S4. Expression of *Oli* RNA in Lin A/15.**

**(A-C)** Lin A/15 in a T2 right hemisegment. **(A)** Max projection of confocal images where Lin A/15 is genetically labeled with myr::GFP (green). **(C1-C2)**. A confocal section of the boxed region in **(A)**. All cells are labeled with dapi (cyan) and *Oli* (red) is labeled with fluorescent probes. Axes: Anterior (A), Medial (L).

**(C)** 3D reconstruction of the segmented Lin A/15 cells on **(A)** from a lateral perspective (**see STAR method)**. Axes: Anterior (A), Ventral (V). Each cell is color-coded based on *Oli* RNA relative expression from low (blue) to high (brown). The *Oli* mRNA spot are represented by spheres (**see STAR method)**.

**(D1-d2)** Plot of the expression level of the *jim* RNA **(D1)** in the Lin A/15 MNs in **(A)** and plot of the average expression **(D2)** level of the *jim* RNA in 6 Lin A/15 samples as a function of their relative distance to the NB. Note: only the 29 motor MNs most distant to NB are represented.

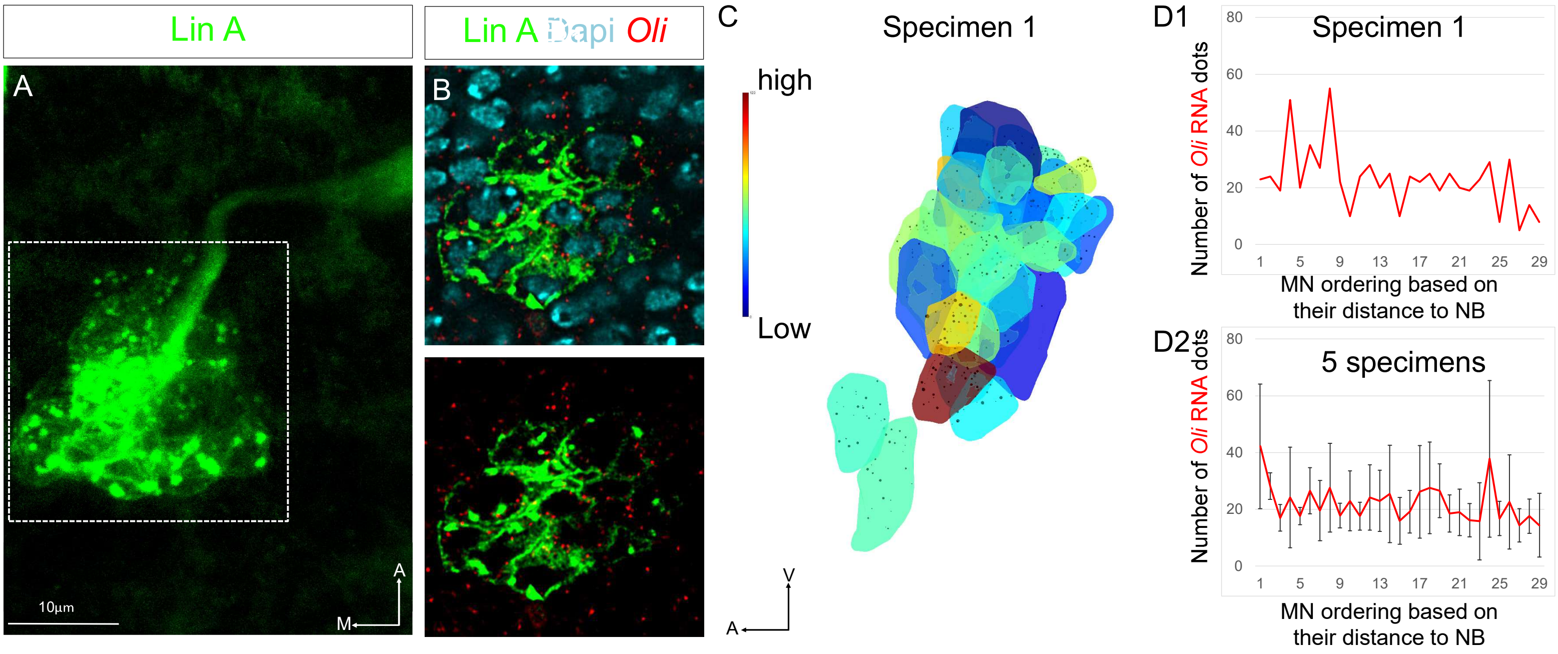
Lin A
A
10µm
A
M
Lin A Dapi Oli
B
C
high
Low
Specimen 1
V
A
D1
Specimen 1
Number of Oli RNA dots
80
60
40
20
0
1
5
9
13
17
21
25
29
MN ordering based on their distance to NB
D2
5 specimens
Number of Oli RNA dots
80
60
40
20
0
1
5
9
13
17
21
25
29
MN ordering based on their distance to NB

**Figure S4**

**Figure S5. Innervation phenotypes in *imp-/-* Lin A/15 clones and numbers MNs in the different gentic conditions.**

**(A1-B3)** Axon targeting phenotypes of *Imp-/-* **(A1-A3)** *Imp-/-*, *tub-P35* **(B1-B3)** and *tub-syp* **(C1-C3)** Lin A/15 MARCM clones. The cuticle is light grey and the axons green; muscle are labeled with *Mhc-RFP* in **(A1-A3)**. The arrow in **(A3)** indicates the lack of a muscle fiber. The arrow in **(A3)** indicates the lack of a muscle innervation in the medial region of the tibia (talm and tadm) N=2/6. The arrow in **(C3)** indicates the lack of a muscle innervation in the medial region of the tibia (talm) N=2/6. Note: **(A1-A3)** Despite a reduced number of MNs produced in *Imp-/-* MARCM clones (Wenyue *et al, unpublished.)* that induces global axonal targeting defects of Lin A/15 iMNs, the distal region of the femur is most affected **(Fig. S5)**

**(C)** Graph of the number of Elav+ neurons observed per Lin A/15 in adult flies under different genetic conditions: *tub>P35*) , *imp-/- tub>P35*, *WT*, *Syp-/-* and tub>Imp.

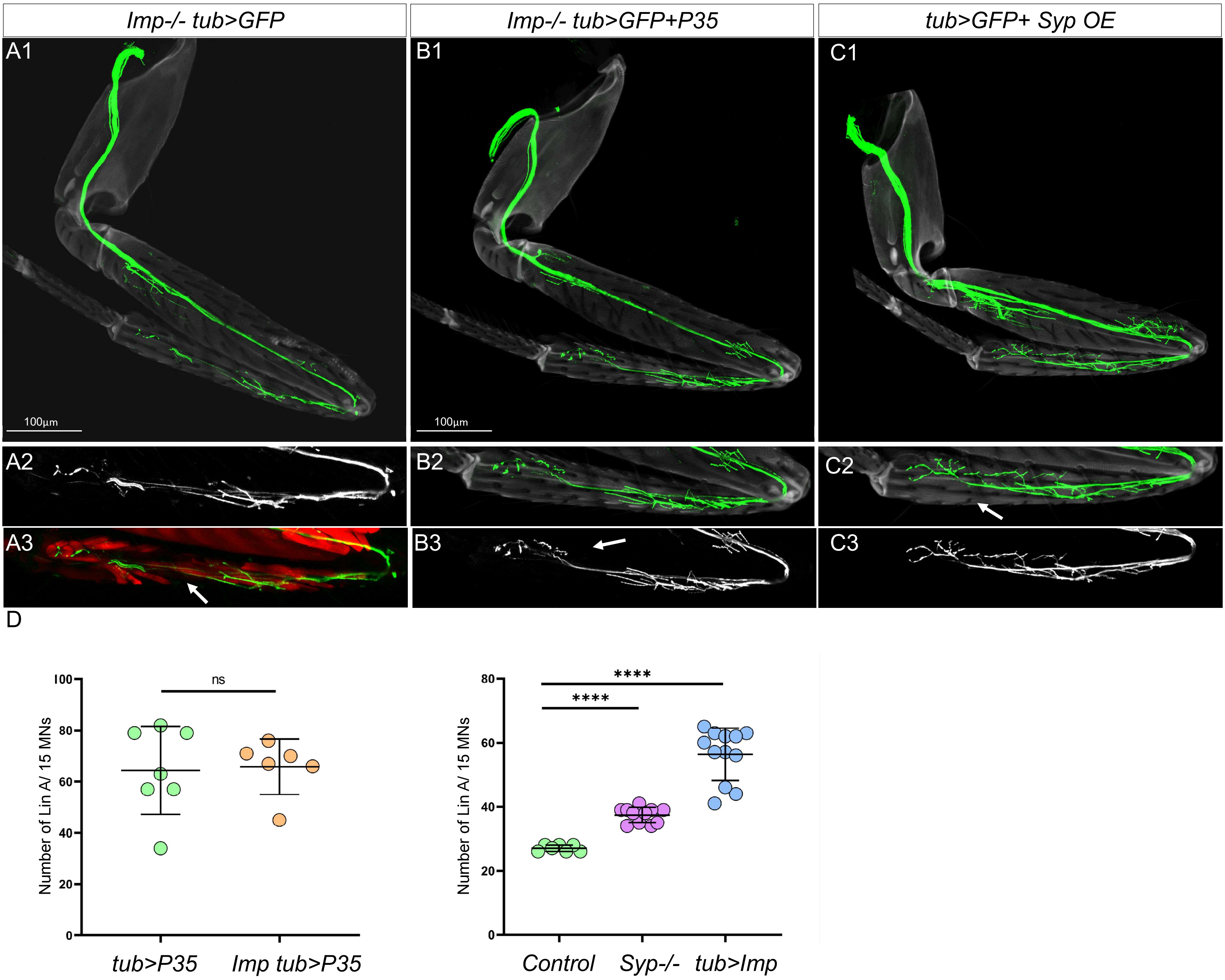

Imp-/- tub>GFP
Imp-/- tub>GFP+P35
tub>GFP+ Syp OE
A1
B1
C1
100µm
100µm
A2
B2
C2
A3
B3
C3
D
ns
Number of Lin A/ 15 MNs
tub>P35
Imp tub>P35
****
****
Number of Lin A/ 15 MNs
Control
Syp-/-
tub>Imp


**Figure S5**

**Figure S6. Imp and Syp do not cross-regulate in L3 VNC**

**(A1-A3, C1-3, F1-F3, H1-H3 )** *WT* **(A1-C4)**, *tub>imp* (**B1-B3, D1-D3, G1-G3**) And *Syp-/-* **(E1-E3, I1-I3)** MARCM clones expressing mCD8::GFP (green) under the control *VGlut-LexA:GAD* **(A1-G3,** tub-Gal4 **(H1-I3)** and immulabaled with anti-Nvy (red) and anti-Br (blue) **(A1-B3)**, anti-Oli (red) **(C1-E3)**, anti-Syp (cyan) **(F1-G3)** and anti-Imp (red) **(H1-I3). (A1-A2, B1-B2, C1-C2, D1-D2, E1-E2, F1-F2, G1-G2, H1-H2, I1-I2)** are confocal sections from ventral to dorsal. **(A3, B3, C3, D3, E3, F3, G3, H3, I3)** Plots of the relative position of each Lin A/15 cell from a lateral perspective; the Lin A/15 cells expressing a giving TF are color-coded in red or blue according the immunostaining in (**A1-A2, B1-B2, C1-C2, D1-D2, E1-E2, F1-F2, G1-G2, H1-H2, I1-I2)**

**(J-N)** Graph of the number of VGlut>GFP+ **(J-L)** or tub>GFP+ **(M-N)** Lin A/15 MNs in *tub>imp vs WT* **(J-K, M),** *Syp-/- vs WT* **(L-N)** expressing Nvy and Broad, **(J)**, Oli **(K-L)**, Imp **(M)**, and Syp **(N)**

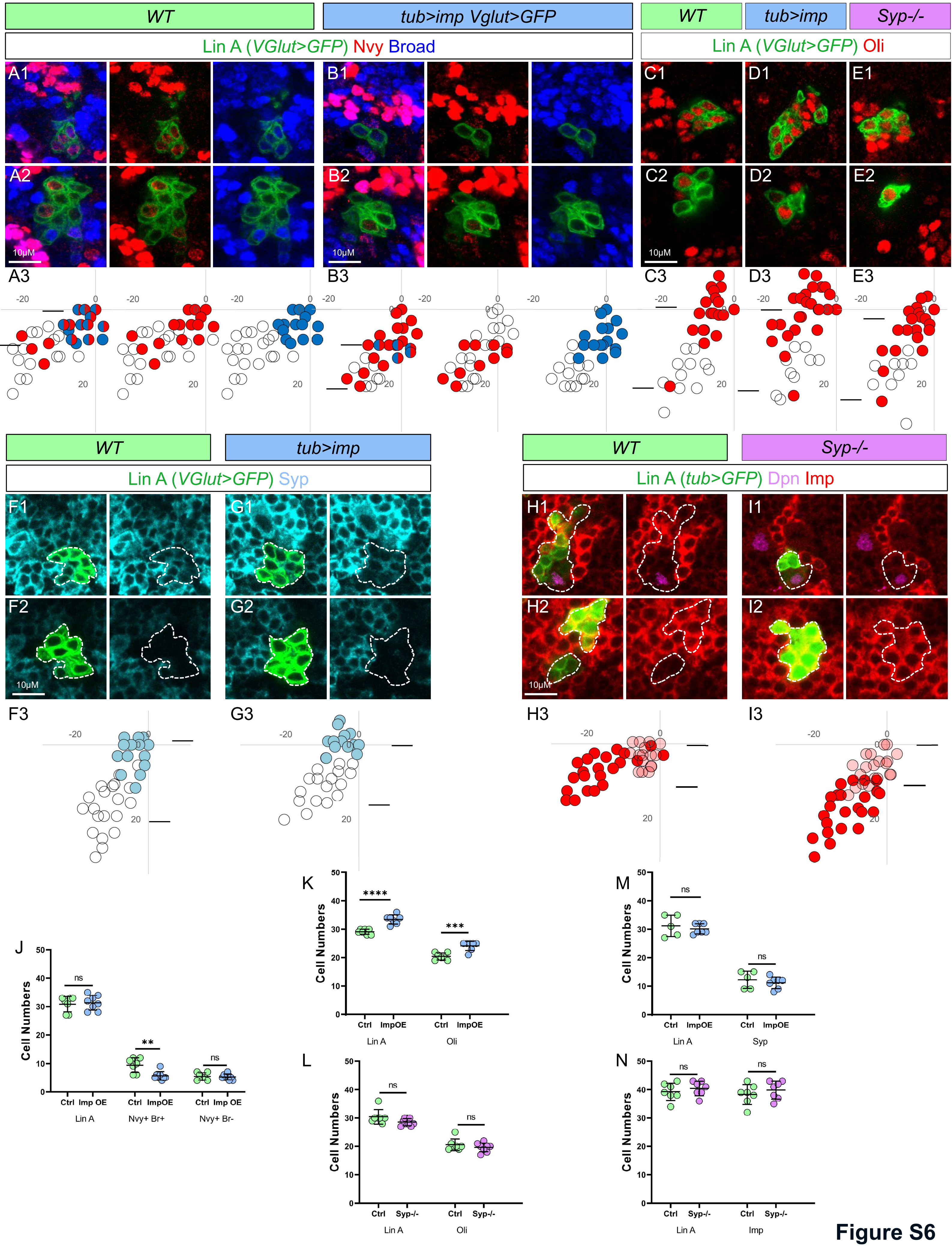
WT
tub>imp Vglut>GFP
WT
tub>imp
Syp-/-
Lin A (VGlut>GFP) Nvy Broad
Lin A (VGlut>GFP) Oli
A1
B1
C1
D1
E1
A2
B2
C2
D2
E2
10μM
A3
B3
C3
D3
E3
WT
tub>imp
WT
Syp-/-
Lin A (VGlut>GFP) Syp
Lin A (tub>GFP) Dpn Imp
F1
G1
H1
I1
F2
G2
H2
I2
F3
G3
H3
I3
J
K
L
M
N
Cell Numbers
Ctrl
Imp OE
ImpOE
Syp-/-
Lin A
Nvy+ Br+
Nvy+ Br-
Oli
Syp
Imp
ns
**
****
***

Figure S6

**Figure S7. Axon targeting phenotypes of *Oli-/-* Lin A/15 MARCM clones**

**(A)** Axon targeting phenotypes of *Oli-/-* Lin A/15 MARCM clones. The cuticle is light grey and the axons green. The arrowhead indicates the lack of a muscle innervation. The arrow in **(A3)** indicates the lack of a muscle innervation in the medial region of the tibia (talm and tadm) N=2/6. The fedm in the trochanter is never targeted (N=13/13) and the tirm in distal femur is less or never innervated (N=9/13). In the distal tibia, the tarm1/2 as well as the distal region of the tadm or talm (N=6/13) are occasionally less or no innervated in *oli-/-* Lin A clones.

*Oli-/- VGlut>GFP*

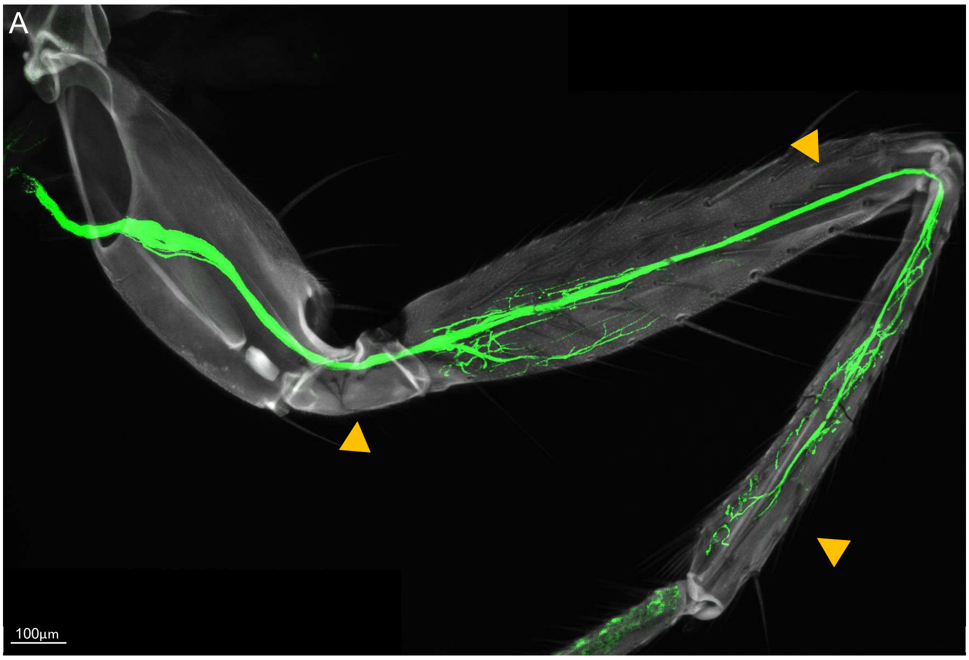


**Figure S7**

**Figure S8. Expression of *imp* and *syp* in iMNs**

**(A1)** Max projection of confocal of the thoracic segments of a L3 VNC containing the six Lin A/15 genetically label with GFP (green).

**(B1, C1, D1)** Confocal sections of the thoracic segments of a L3 VNC in **(A).** All cells are labeled with dapi (blue); *Syp* (purple) and *Imp* (red*)* nascent mRNAs are labeled with intronic fluorescent probes.

**(A2, B2, C2, D2)** magnification of the doted box in **(A1, B1, C1, D1)**

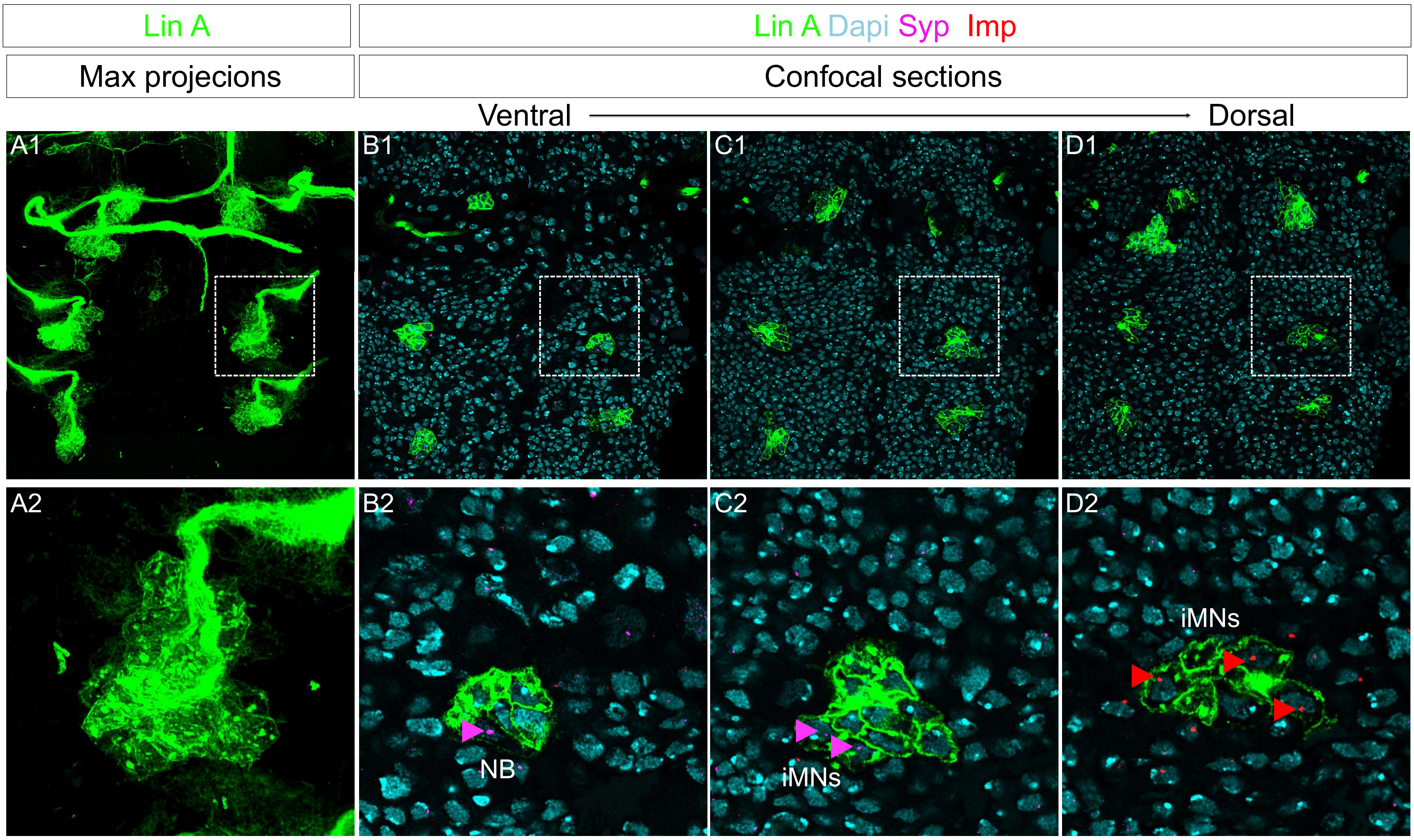
Lin A
Lin A Dapi Syp Imp
Max projecions
Confocal sections
Ventral
Dorsal
A1
B1
C1
D1
A2
B2
C2
D2
NB
iMNs
iMNs

**Figure S8**

### Supplemental notes

#### Supplemental note S1. Lin A/15 MARCM clones vs Lin A/15 memory system

The one-spot MARCM technique makes it possible to visualize the progeny of only one of the daughter cells derived from a common progenitor (Lee and Luo, 2001). Consequently, the generation of an NB MARCM Lin A/15 clone at the larval stage, when the NB is quiescent, makes it possible to visualize the first MN born or the next 28 MNs, but not all together. However, the Lin A/15 memory system (Awasaki et al., 2014; Lactin and Truman., 2016) allows the visualization of all 29 MNs.

#### Supplemental note S2. Link between birth order and muscle targets: Corrections of Myungin paper

A previous study revealed a link between the Lin A MN birth order and the muscles they target (Baek and Mann, 2009). In our study **(Figure 1)**, we made two corrections by adding two MNs (Lin A Tr1 and B1), not described in (Baek and Mann, 2009). The relative birth orders of the 28 Lin A MNs have been characterized by inducing GMCs MARCM clones at different time points during larval stages. The generation of GMC MARCM clones has allowed the determination of the relative birth order of 27 MNs out the 28 MNs. In our study, when we generated an NB MARCM clone to label the 28 Lin A MNs, we always marked an MN targeting the trochanter segment, an MN not considered to be part of Lin A (Baek and Mann, 2009). We named this MN, Lin A Tr1. Based on the shape of the axonal terminal branches, we propose that this MN has been misidentified (Baek and Mann, 2009) with Lin G, a lineage supposedly producing a single MN. Here, we propose that Lin G is not a lineage by itself (discovered by generating an NB MARCM clone) but a GCM MARCM clone. Moreover, we conclude that Lin A Tr1 MN is the first born MN of the 28 MNs by comparing in the Baek and Mann article (2009), the last time point where the Tr1 GMC MARCM clone can be induced with the other GMC MARCM clones. Finally, the NB one-spot MARCM technique, cannot label the first Lin A MN. The twin spot QMARCM/MARCM has revealed that the first-born MN from Lin A targets a body wall muscle (Enriquez et al., 2018). Here, we name this MN: B1.

### Supplemental experimental procedures

#### Experimental procedures S1. Testing the model by co-staining

The cell cluster detection method predicts the combinations of TFs expressed in each immature MN. To validate the positive-cell cluster detection method and to determine its accuracy, we tested its predictions by performing co-stainings. Among 256 possible co-stainings, we chose N=12 co-staining combinations, serving as a proof-of-concept. We chose those combinations to test two parameters of the positive-cell cluster detection method: the accuracy of the position of the positive-cell cluster and the accuracy of the number of cells per cluster when 2 cell clusters were detected (length of the positive cell cluster).

*The accuracy of the position of the positive cell cluster*

The number of cells expressing a given TF (length of the positive-cell cluster) is not a prediction of the model when only one cluster is detected because this parameter is based on real data: average number of positive cells expressing a given TF from all our experiments. However, we tested the accuracy of the position on the x' axis of the positive positive-cell cluster by performing co-staining between TFs that have partially overlapping cluster coverage.

**Kr and Zfh2 co-staining. Prediction of the PCCD**: The model predicts 3 to 5 cells expressing both TFs with a higher probability than only 3 cells because there is a higher chance for cell #5 rather than cell #1 to express Kr (coverage index at cell #5 for Kr is 0.9 and 0.4 at cell #1) and low chance for cell #5 to express Zfh2 (coverage index at cell #5 for Zfh2 is 0.3). **Result of the co-staining**: Number of cells expressing Kr and Zfh2=3. We concluded, as predicted by the model, that cell #5 expresses Kr whereas cell #1 did not, and cell #5 is Zfh2 negative.

**Lov and Zfh2 co-staining. Prediction of the PCCD:** The model predicts 1 to 3 cells expressing both TFs with a higher probability than 2 cells because there is higher chance for cell #3 rather than cell #7 to express Lov (coverage index at cell #3 for Kr is 0.6 and 0.4 at cell #7) and low chance for cell #5 to express Zfh2 (coverage index at cell #5 for Zfh2 is 0.3). **Result of the co-staining**: Number of cells expressing Lov and Zfh2=2. Lov antibody seems to be sensitive to fixation conditions and we found one more cell expressing low level of Lov after optimizing the fixation conditions **(Fig S2)**. Consequently, we corrected the length of the positive cell cluster of Lov from 4 to 5 and included both cell #3 and cell #7. We included cell #3 as the cell expressing low level of Lov because the cell expressing a low level of Lov is Zfh2+.

**Chinmo and Jim co-staining. Prediction of the PCCD:** The model predicts to have 3 to 5 cells expressing both TFs. **Result of the co-staining**: Number of cells expressing Chinmo and Jim=4. Here, we have two possibilities: cell #15 Jim+ and cell #19 Chinmo- or cell #15 Jim- and cell #19 Chinmo+. In the PPCD method the length of the positive cell cluster is 18.5. After including the Chinmo+ cells from co-staining experiments the average number of cell expressing Chinmo is closer to 19. This variation between samples is due to the weak expression of Chinmo in one Jim+ cell localized ventrally, which is sometimes not detected **(Fig S2)**. We concluded that cell #23 expresses Jim while cell #15 does not.

**Conclusion:** the combinatorial expression of TFs predicted by the method seems to be accurate since the co-staining validated the predictions.

*The number of cells per cluster when 2 cell clusters are detected (length of positive cell cluster ) and accuracy of the position of the positive cell cluster.*

We tested the length of the positive dorsal cluster (left cluster) when two clusters were detected, by performing co-staining between TFs that were predicted to be completely overlapping in the dorsal cluster and not the ventral one. We then tested the accuracy of the position of the positive cell cluster as described in the previous paragraph. We tested the length, as well as the position, of cell clusters expressing **RunxA, Zfh1, Oli and Nvy.**

**RunxA and Chinmo co-staining** (testing the length of the dorsal RunxA+ cluster)**. Prediction of the PCCD**: The model predicts a length of the dorsal RunxA+ cluster = 5.1 and a number of cells expressing RunXA and Chinmo= 5. **Result of the co-staining:** Number of cells expressing Zfh1 and Chinmo= 5. We concluded that the number of cells expressing RunxA in the dorsal is cluster is correct.

**RunxA and Jim co-staining** (testing the position of the dorsal RunxA+ cluster)**. Prediction of the PCCD**: The model predict 0 to 1 cell expressing both. **Result of the co-staining:** Number of cells expressing RunxA and Jim =0. This result indicates that cell #15 is not RunxA+, Jim+. Because we demonstrated in the previous paragraph that cell #15 is Jim-, we could not conclude if the cell #10 or the cell #15 is RunxA+.

**Zfh1 and Chinmo co-staining** (testing the length of the dorsal Zfh1+ cluster)**. Prediction of the PCCD**: The model predicts a length of the dorsal Zfh1+ cluster= 10.8 and a number of cells expressing Zfh1 and Chinmo= 10 or 11. **Result of the co-staining:** Number of cells expressing Zfh1 and Chinmo=12. Based on these results, we made a correction of the length of the dorsal Zfh1+ cluster from 10.8 to 12.

**Zfh1 and Zfh2 co-staining** (testing the position of the dorsal Zfh1+ cluster)**. Prediction of the PCCD**: The model predicted 2 to 4 cells expressing both TFs. **Result of the co-staining:** Number of cells expressing Zfh1 and Zfh2=2. We concluded that cell #2 does not express Zfh1 and cell #5 does not express Zfh2

**Zfh1 and Kr co-staining** (testing the position of the dorsal Zfh1+ cluster)**. Prediction of the PCCD**: The model predicts 2 to 4 cells expressing both TFs. **Result of the co-staining:** Number of cells expressing Zfh1 and Kr =3. We had concluded that cell #5 is Kr+, confirming that cell #2 does not express Zfh1.

Based on these three co-stainings including Zfh1, we concluded that cell #13 expresses Zfh1 while cell #2 does not. Moreover, we made a correction by adding cell #14 into the Zfh1+ cluster to respect a length of the Zfh1+ cluster =12.

**Oli and Zfh2 co-staining** (testing the length of the dorsal Oli+ cluster)**. Prediction of the PCCD**: The model predicts a length of the dorsal Oli+ cluster = 2.8 and a number of cells expressing Oli and Zfh2=2 or 3. **Result of the co-staining:** Number of cells expressing Zfh1 and Kr =2. Based on these results we made a correction of the length of the of Oli+ cell cluster for the first Oli cluster. The length of the Oli+ cell cluster =2 instead of 2.8.

**Oli and Kr co-staining** (testing the position of the dorsal Oli+ cluster)**. Prediction of the PCCD**: The model predicts to have 1 to 3 cells expressing both. **Result of the co-staining**: Number of cells expressing Oli and Kr=1. We concluded that cell #1 expresses Oli whereas cell #3 does not.

**Nvy and Chinmo** (testing the length of the dorsal Nvy+ cluster)**. Prediction of the PCCD**: The model predicts a length of the dorsal Nvy+ cluster= 3.6 and a number of cells expressing Nvy and Chinmo= 3 or 4. **Result of the co-staining:** Number of cells expressing Nvy and Chinmo=4. Notably , we notice that the dorsal Nvy+ cluster express higher level of nvy compare to the ventral cluster.

**Nvy and Jim** (testing the position of the dorsal Nvy+ cluster)**. Prediction of the PCCD:** The model predicts 0 to 2 cells expressing both TFs. **Result of the co-staining**: Number of cells expressing Nvy and Jim= 2. However, these two cells express a very low level of Nvy and are located ventrally in the 14 samples analyzed. We concluded that cell #12 instead of #16 is Nvy+. See Discussion for comments on the ventral clusters and why the PCCD method did not predict that 2 cells of the ventral Nvy+ cluster express Jim.

**Conclusion:** The length of the cluster coverage when two clusters are detected, is accurate. No corrections or sometimes corrections of one cell to cluster length were made.

**Note for the ventral clusters:** The PCCP method probably underestimated the number of cells that should be included on the left region of clusters located near the NB. Indeed, the frequency of TF expression in a cell close to the NB was low, not because of the variation in the cell positioning between the samples, but because the cell may not have been born in some samples. This would artifactually reduce the frequency of TF expression at positions close to the NB. This bias could be overcome in a future version of the PCCD method. However, the Nvy and Jim co-staining revealed that the errors were small. Finally, all ventral clusters had a gradient expression from high (ventral) to low (dorsal). Consequently, the boundary of the left region of these clusters was not sharp due to weak TF expression that was sometimes difficult to detect.

**Experimental procedures S2. The pipeline for smFISH analysis**

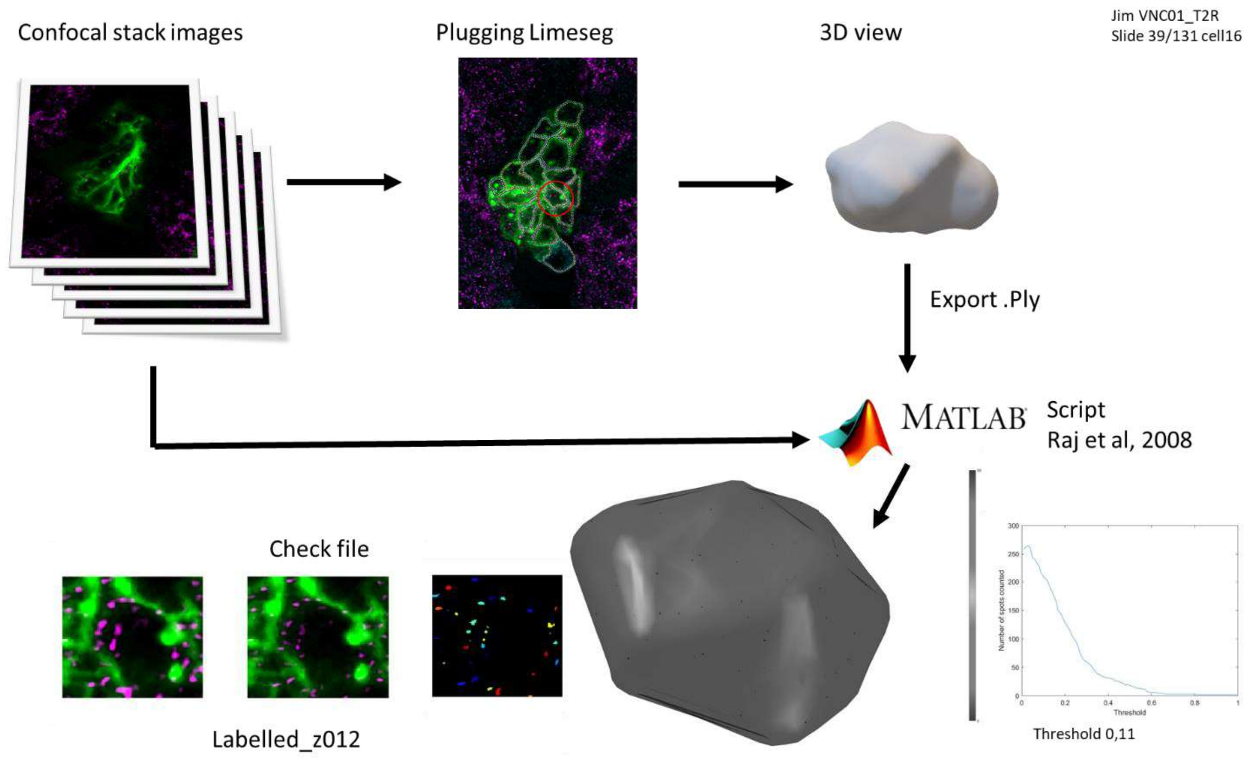
